# Covariate Assisted Principal Regression for Covariance Matrix Outcomes

**DOI:** 10.1101/425033

**Authors:** Yi Zhao, Bingkai Wang, Stewart H. Mostofsky, Brian S. Caffo, Xi Luo

**Author notes:** September 13, 2018.

## Abstract

Modeling variances in data has been an important topic in many fields, including in financial and neuroimaging analysis. We consider the problem of regressing covariance matrices on a vector covariates, collected from each observational unit. The main aim is to uncover the variation in the covariance matrices across units that are explained by the covariates. This paper introduces *Covariate Assisted Principal* (CAP) regression, an optimization-based method for identifying the components predicted by (generalized) linear models of the covariates. We develop computationally efficient algorithms to jointly search the projection directions and regression coefficients, and we establish the asymptotic properties. Using extensive simulation studies, our method shows higher accuracy and robustness in coefficient estimation than competing methods. Applied to a resting-state functional magnetic resonance imaging study, our approach identifies the human brain network changes associated with age and sex.

## 1 Introduction

Modeling variances is an important topic in the statistics and financial literature. In linear regression with heterogeneous errors, various (generalized) linear models have been proposed to model the error variances using the covariates directly or indirectly as a function of the mean (see for example Box and Cox (1964); Carroll et al. (1982); Smyth (1989); Cohen et al. (1993)). These models use separate regression models of the covariates to predict a scalar variance parameter of the error, as well as the mean of the response. Usually, the goal is to improve the efficiency of estimating the mean regression model, while the variance regression model is of less interest.

Regression models for covariance matrices were studied before under different settings. For time series, the “autoregressive conditionally heteroscedastic” (ARCH) models (Engle and Kroner, 1995) were developed to model temporal heteroscedasticity. Anderson (1973) proposed an asymptotically efficient estimator for a class of covariance matrices, where the covariance matrix is modeled as a linear combination of symmetric matrices. Chiu et al. (1996) proposed to model the elements of the logarithm of the covariance matrix as a linear function of the covariates. Pourahmadi (1999) considered another type of matrix decomposition, where the covariates predict linearly the unconstrained elements in the Cholesky decomposition. However, this approach is not order invariant, and requires the matrix columns/rows follow a meaningful ordering. These matrix regression models usually require a large number of parameters to be estimated.

Several approaches were proposed to extend matrix outcome regression models to high dimensions. Hoff and Niu (2012) introduced a regression model, where the covariance matrix is a parsimonious quadratic function of the explanatory variables. Applying low-rank approximation techniques, Fox and Dunson (2015) generalized the framework to a nonparametric covariance regression model and enabled scaling up to high dimensions. In a recent paper, Zou et al. (2017) linked the matrix outcome to a linear combination of similarity matrices of covariates, and studied the asymptotic properties of various estimators under this model. These approaches again model the whole covariance matrix as outcomes, and thus the interpretation could be challenging for large matrices.

Closely related to covariance matrices, principal component analysis (PCA) and related methods are widely used to generate interpretable results for large dimensional data. These methods have been extended to model multiple covariance matrices. Flury (1984) and Flury (1988) introduced a class of models, called common principal components models, to uncover the shared covariance structures. Boik (2002) generalized these models using spectral decompositions. Hoff (2009) developed a Bayesian hierarchical model and estimation procedure to study the heterogeneity in both the eigenvectors and eigenvalues of covariance matrices. Assuming that the eigenvectors span the same subspace, Franks and Hoff (2016) extended this to the so-called high dimensional setting with large *p* and small *n*. It is unclear, however, how these methods can be extended to incorporate multiple covariates.

In the application area of neuroimaging analysis, PCA-type methods are becoming increasingly popular for modeling covariance matrices, partly because of their desirable interpretability and computational capability for analyzing large and multilevel observations. Covariance matrices (or correlation matrices after standardization) of multiple brain regions are also commonly known as functional connectivity analysis (Friston, 2011). Decomposing the covariance matrices into separate components enable identification of coherently active brain subnetworks (Poldrack et al., 2011), and usually a few principal components are needed to explain the variation in neuroimaging data (Friston et al., 1993). As before, it is unclear how these methods can be extended to include multiple covariates.

Indeed, modeling the covariate-related alterations in covariance matrices is an important topic in neuroimaging analysis, because changes in functional connectivity have been found to be associated with various demographic and clinical factors, such as age, gender, and cognitive behavioral functions including developmental and mental health capacities (Just et al., 2006; Wang et al., 2007; Luo et al., 2011; Mennes et al., 2012; Hafkemeijer et al., 2015; Park et al., 2016). A commonly implemented method to analyze the covariance changes is to regress one matrix entry on the covariates, and this model is repeatedly fitted for each matrix element (see, for example, Wang et al. (2007) and Lewis et al. (2009)). Though this approach has good interpretability and is scalable, it suffers from the multiplicity issues, because of the large number of regressions involved. For example, *p*(*p* − 1)*/*2 regressions for *p* brain regions. Adapting the covariance regression model proposed in Hoff and Niu (2012), Seiler and Holmes (2017) introduced a simplified model to analyze a large and multilevel neuroimaging dataset.

In this paper, we propose a *Covariate Assisted Principal (CAP) regression* model for multiple covariance matrix outcomes. This model integrates the PCA principle with a generalized linear model of multiple covariates. Analogous to PCA, our model aims to identify linear projections to allow for interpretability of the covariance matrices, while being computationally feasible for large data. Unlike PCA, our method targets the projections that are associated with the covariates. This enables us to study the changes in covariance matrices associated with subject-specific factors, such as individual demographic or disease information.

This paper is organized as follows. In Section 2, we introduce our proposed CAP regression model. Section 3 presents the estimation and computation algorithms in identifying the proposed principal projection directions. We compare the performance of our proposed methods with competing approaches through simulation studies in Section 4. We then apply our methods to a real fMRI dataset in Section 5. Section 6 summarizes the paper with a summary and discussion of future directions.

## 2 Model

For each *i* ∈ {1, …, *n*}, let **y**_*it*_ ∈ ℝ^*p*^, *t* = 1, …, *T_i_*, be independent and identically distributed random samples from a multivariate normal distribution with mean zero and covariance matrix, Σ_*i*_, where Σ_*i*_ may depend on explanatory variables, **x**_*i*_ ∈ ℝ^*q*−1^. In our application example, **y**_*it*_ is a sample of brain fMRI measurements of *p* regions, and **x**_*i*_ is a vector of covariates postulated to be related to fMRI measurements, both collected from subject *i*. We assume that there exists a vector ***γ*** ∈ ℝ^*p*^ such that
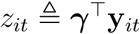
satisfies the following multiplicative heteroscedasticity model:

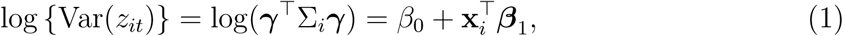

where *β*_0_ and ***β***_1_ are model coefficients. The logarithmic linear model follows from Harvey (1976) in which Σ_*i*_ is a scalar.

A toy example of this model (*p* = 2) is shown in Figure 1. The covariance matrices, represented by the contour plot ellipses, vary as the covariate *x* varies. On the first projection direction (PD1) with the largest variability, there is no variation under different *x* values. However, the variance in the second direction (PD2) decreases as *x* increases. Our proposed model (1) thus aims to identify the second projection direction. In other words, the objective is to discover the rotation such that the data variation in the new space can be best characterized by the explanatory variables.

**Figure 1:**
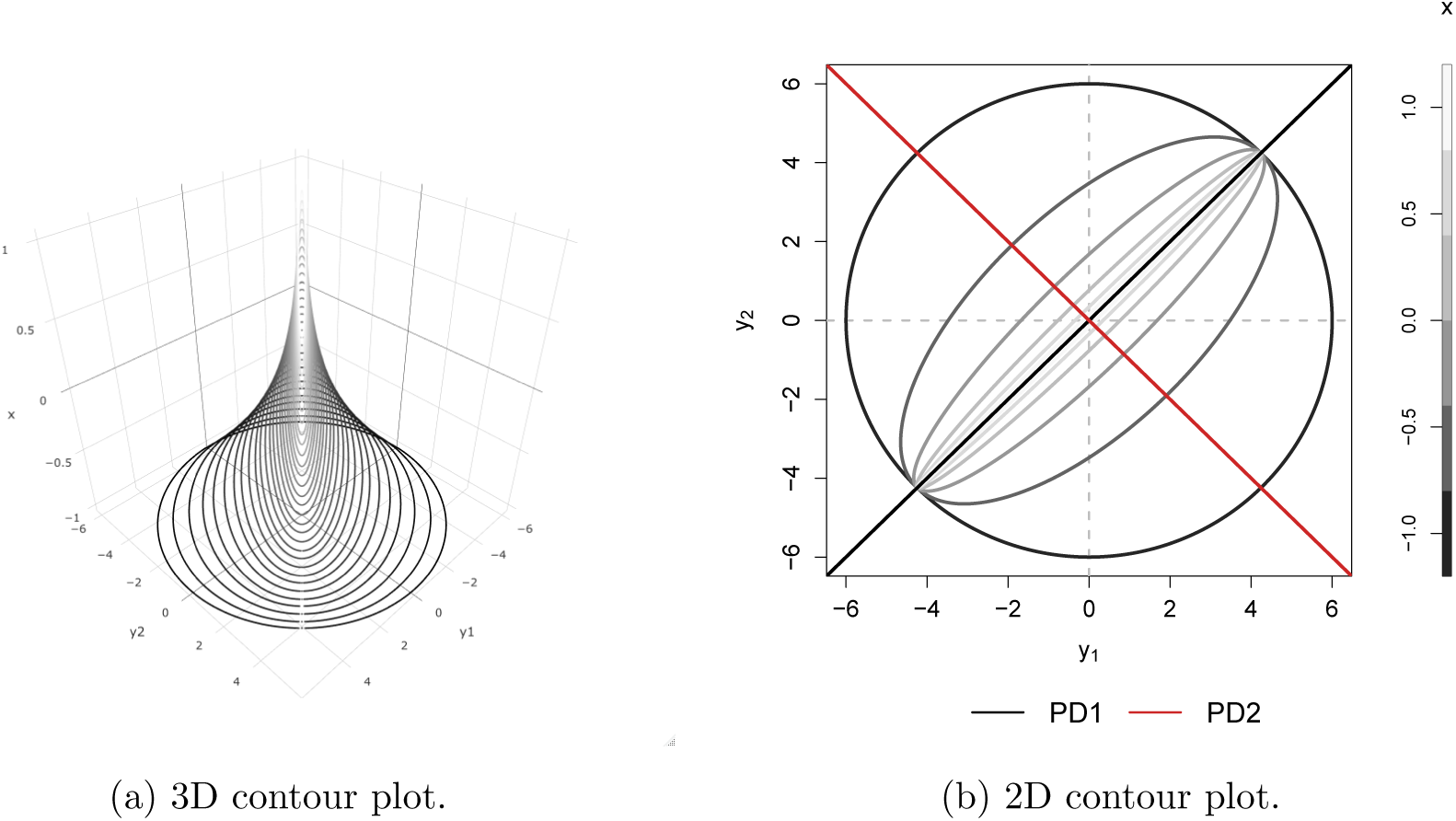
Covariance matrices, shown as contour plot ellipses when *p* = 2, vary as a continuous *X* varies (*z*-axis in (a) and gray/color scales in (b)).

Compared with existing methods, our proposed model has two main advantages. First, different from the model proposed by Hoff and Niu (2012) and Zou et al. (2017), which directly model Σ_*i*_ by linear combinations of symmetric matrices constructed out of **x**_*i*_, we assume a log-linear model for the variance component after rotation. The linear form allows easy interpretation of the regression coefficient and provides the modeling flexibility shared by all other (generalized) linear models, such as interactions. The projection enables computational scalability similar to PCA. The common principal component approach, studied in Flury (1984), only allows the eigenvalues to vary across a group indicator, our model (1) provides a direct model of multiple covariates, including continuous ones. This enables studying the covariate-related changes in covariances in our fMRI experiment. Second, our model relaxes the standard complete common principal component assumption imposed in Flury (1984) and Boik (2002), and we assume that there exists at least one projection direction such that model (1) is satisfied. This partial common diagonalization assumption is more realistic for data with higher dimensions.

## 3 Method

We propose to estimate the model parameters by maximizing the likelihood function under a quadratic constraint:

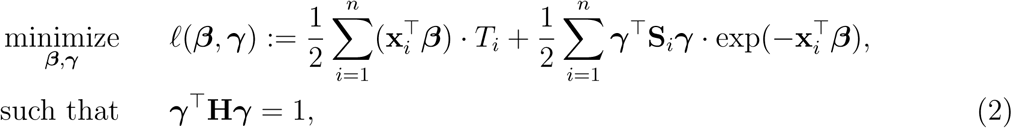

where *ℓ*(***β***, ***γ***) is the negative log-likelihood function (ignoring constants),
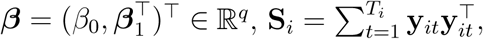
, and **H** is a positive definite matrix in ℝ^*p*×*p*^. Without the constraint, *ℓ*(***β***, ***γ***), for any fixed ***β***, is minimized by ***γ*** = **0**. Thus the constraint is critical.

Two natural choices of **H** in the constraint are:

(C1) **H** = **I** which is equivalent to a unit constraint under *ℓ*_2_-norm, i.e.,

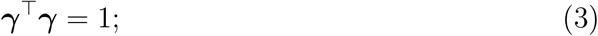

(C2) **H** =
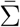
which is equivalent to a unit constraint with respect to the average sample covariance, i.e.,

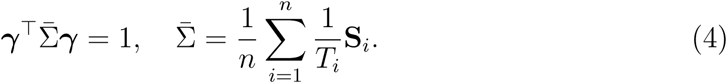

(C1) is inspired by standard PCA. The second one is by common principal component analysis. We show in the following proposition that (C1) will lead to a solution that is less appealing in certain situations.

### Proposition 1.

*When* **H** = **I** *in the optimization problem* (2), *for any fixed **β***, *the solution of **γ** is the eigenvector corresponding to the minimum eigenvalue of matrix*

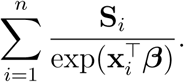

The matrix
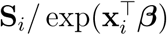
can be regarded as a normalization on the covariance matrices based on the explanatory variables. Thus, constraint (C1) achieves the projection direction with the lowest normalized data variation. In the Appendix Section D, we further discuss the property of these two constraints using examples. We will focus on constraint (C2) in this paper because the signals are usually not associated with the smallest eigenvalue in most scenarios.

### 3.1 Algorithm

The optimization problem (2) is biconvex. We propose to solve the optimization problem by block coordinate descent. For given ***γ***, the update of ***β*** is obtained by the Newton-Raphson algorithm. For given ***β***, the solving for ***γ*** requires quadratic programming. Though generic quadratic programming packages could be used, we derive the explicit solution in Proposition A.1 in the supplementary material. The algorithm is summarized in Algorithm 1. This algorithm works for any positive definite **H**. To obtain robustness against obtaining a solution in a local minimum, we propose to randomly choose a series of initial values and take the estimate with the lowest objective function value.

### 3.2 Extension for finding multiple projection directions

It is possible that more than one projection direction is associated with the covariates. We propose to find these directions sequentially. This is modified from the strategy of finding multiple principal components one by one.

#### Algorithm 1 A block coordinate descent algorithm for solving optimization problem (2).

**Input:**

**Y**: a list of data where the ith element is a *T_i_* × *p* data matrix

**X**: *a n* × *q* matrix of covariate variables with the first column of ones *β*^(0)^, *γ*^(0)^ : initial values

**Output**:
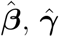

Given (*β*^(*s*)^, *γ*^(*s*)^ from the sth step, for the (*s* + 1)th step:

(i) update *β* by

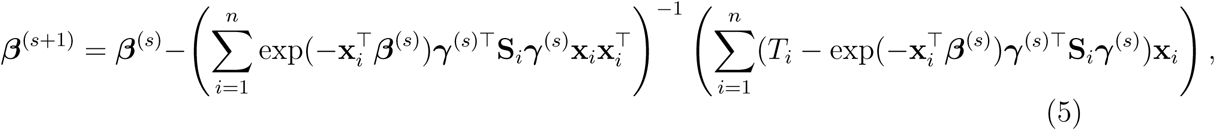

where
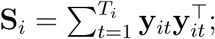
(ii)update *γ* by solving

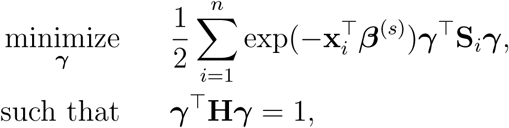

where **H** = **I** under (C1) and
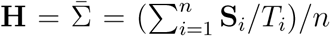
under (C2), using Proposition A.1. Repeat steps (i)-(ii) until convergence.

Suppose Γ^(*k*−1)^ = (***γ***^(1)^, …, ***γ***^(*k*−1)^) contains the first (*k* − 1) components (for *k* ≥ 2), and let
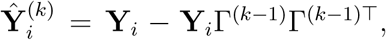
where **Y**_*i*_ = (**y**_*i*1_, …, **y**_*iT_i_*_)^⊤^ for *i* = 1, …, *n*. We cannot directly apply Algorithm 1 to
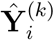
as in PCA algorithms, since
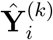
is not of full rank. We introduce a rank-completion step. The whole algorithm is summarized in Algorithm 2. In the algorithm, step (iii) completes the data to full rank by adding nonzero positive eigenvalues to those zero eigencomponents, which are the exponential of model intercept of the corresponding directions. This step also guarantees that there are no identical eigenvalues in the covariance matrix of **Ỹ**_*i*_, which is a necessary condition for unique eigenvector identification.

Analogous to the PCA approach, step (iv) is an orthogonal constraint to ensure that the *k*th direction is orthogonal to the previous ones, which is equivalent to the following optimization problem, for *k* ≥ 2,

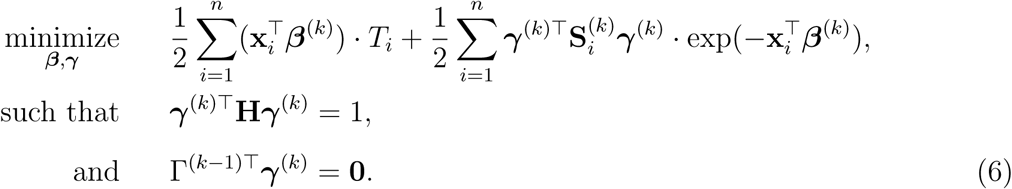

For any fixed ***β***^(*k*)^, we derive an explicit formula for solving ***γ***^(*k*)^, see Section A.2 of the supplementary material. The proof is adapted from Rao (1964, 1973).

### 3.3 Choosing the number of projection directions

We propose a data-driven approach to choose the number of projection directions. Extending the common principal component model, Flury and Gautschi (1986) introduced a metric to quantify the “deviation from diagonality”. Suppose **A** is a positive definite symmetric matrix, the “deviation from diagonality” is defined as

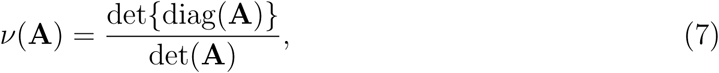

#### Algorithm 2 An algorithm for finding the *k*th projection direction under constraint (C2).

**Input:**

**Y**: a list of data where the *i*th element is a *T_i_* × *p* data matrix

**X**: *a n* × *q* matrix of covariate variables with the first column of ones Γ^(*k*−1)^: a *p* × (*k* − 1) matrix contains the first (*k* − 1) directions

**B**^(*k*−1)^: a *q* × (*k* − 1) matrix contains the model coefficients of the first (*k* − 1) directions

**Output**:
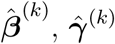

(i) For *i* = 1, …, *n*, let
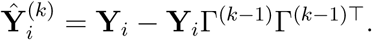
(ii) Apply singular value decomposition (SVD) on
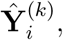
such that
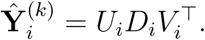
(iii) Let
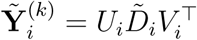
with

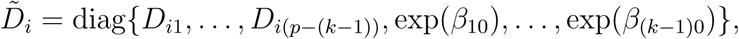

where {*D*_*i*1_, …, *D*_*i*(*p*−(*k*−1))_} are the first (*p* − (*k* − 1)) diagonal elements of matrix *D_i_*, and *β*_10_, …, *β*_(*k*−1)0_ are the intercept of the first (*k* − 1) directions (first row of **B**^(*k*−1)^).
(iv) Treat
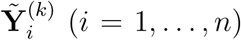
as the new data, and apply Algorithm 1 under constraint (C2) with an additional orthogonal constraint

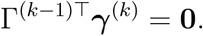

where diag(**A**) is a diagonal matrix with the diagonal elements the same as matrix **A**, and det(**A**) is the determinant of matrix **A**. From Hadamard’s inequality, we have that *ν*(**A**) ≥ 1, where equality is achieved if and only if **A** is a diagonal matrix.

To adapt this metric in our model, we let Γ^(*k*)^ ∈ ℝ^*p*×*k*^ denote the matrix containing the first *k* projection directions. We define the average deviation from diagonality as

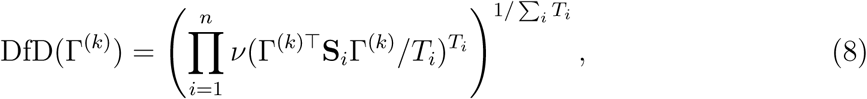

which is the weighted geometric mean of each subject’s deviation from diagonality. As *k* increases, the requirement for Γ(*k*)^*⊤*^**S**_*i*_Γ^(*k*)^ to be a diagonal matrix, as in Flury and Gautschi (1986), may become more stringent. In practice, we can plot the average deviation from diagonality and choose the first few projection directions with DfD value close to one or choose a suitable number right before a sudden jump in the plot. See an example in Section E of the supplementary material.

### 3.4 Analysis under a Common Principal Component Model

We need additional assumptions to perform theoretical analysis. Following Flury (1986), we assume that the covariance matrices Σ_1_, …, Σ_*n*_ can be diagonalized by the same orthogonal matrix. That is, there exists an orthogonal matrix Γ, such that

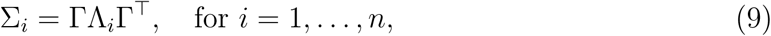

where Γ = (***γ***_1_, …, ***γ**_p_*) and Λ_*i*_ = diag{λ_*i*1_, …, *λ_ip_*}. Suppose the eigenvalues are ordered as
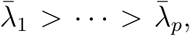 where
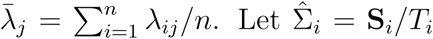 denote the sample covariance matrix. Suppose Φ = (***ϕ***_1_, …, ***ϕ***_*p*_) and Δ_*i*_ = diag{*δ*_*i*1_, …, *δ_ip_*} are the maximum likelihood estimator of Γ and Λ_*i*_ (*i* = 1, …, *n*), respectively, using the method proposed in Flury (1984). Flury (1986) showed that they are both consistent estimators, and thus
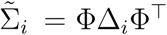 is a consistent estimator of Σ_*i*_. Therefore, we have

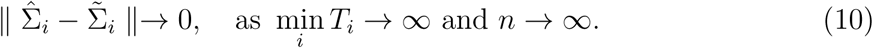

Based on (10), we replace **S**_*i*_ by the consistent estimator
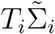 in our optimization problem (2). Since Φ is the orthonormal eigenbasis, ***γ*** can be represented by the linear combination of the columns in Φ, i.e.,
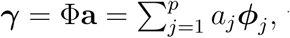 where **a** = (*a*_1_, …, *a_p_*)^*⊤*^. The optimization problem (2) is reformulated as:

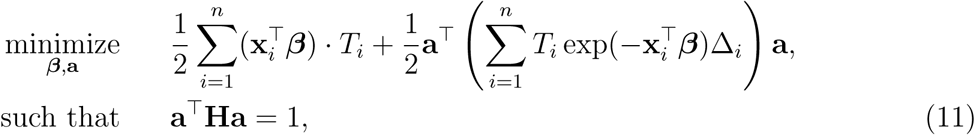

where **H** = **I** under (C1) and
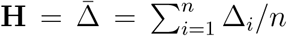 under (C2). With given ***β***, under constraint (C1), it is equivalent to solve

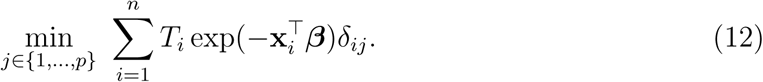

Suppose the eigenvectors are ordered based on the average eigenvalues
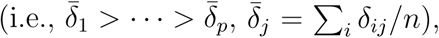 we have **â** = ***ϕ**_p_*. Therefore, under constraint (C1), the method yields the common eigenvector with the lowest average eigenvalue. Now consider the constraint (C2). Let **b** = (*b*_1_, …, *b_p_*)^*⊤*^ with
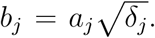 Minimizing the objective function in (11) under constraint (C2) is equivalent to solving the following problem,

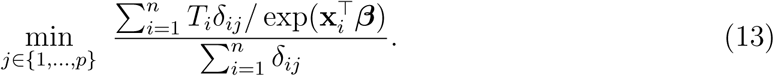

Suppose
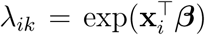 satisfies the model assumption, with *T_i_* = *T*, the minimizer of above optimization problem is *k* if

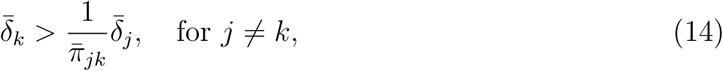

where
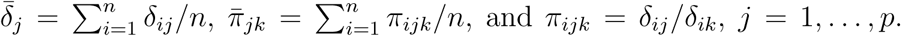 Since *δ_ij_* is a consistent estimator of λ_*ij*_ for *i* = 1, …, *n* and *j* = 1, …, *p*, we impose the following condition.

#### Condition 1

(Eigenvalue condition). *Assume* Σ_*i*_ = ΓΛ_*i*_Γ^⊤^ *is the eigendecomposition of* Σ_*i*_ *with* Γ = (***γ***_1_, …, ***γ**_p_*) *an orthogonal matrix and* Λ_*i*_ = diag{λ_*i*1_, …, λ_*ip*_} *a diagonal matrix*, *for i* = 1, …, *n. The eigenvalues are ordered as*
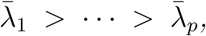 *where* 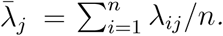 *Suppose there exists k* ∈ {1, …, *p*} *such that λ_ik_* =
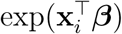 *satisfies the model assumption*, *and assume*

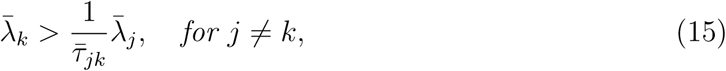

*where*
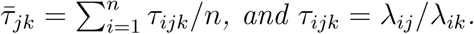

Under this condition, we propose a min-max algorithm (Algorithm 3) to identify the common principal component with eigenvalues that fit the log-regression model (1) and meanwhile explain large variations in the data. We call this algorithm a min-max approach as it contains a minimization (of the objective function) and maximization (of data variation) steps. To acquire the first *k* (*k* ≥ 2) directions, we propose to order those *p_s_* components that satisfy *ŝ*(*j*) = *j* in step (iv) by the average eigenvalues and return the first min{*k*, *p_s_*} components. Thus this algorithm also provides an estimate of the number of components.

#### 3.4.1 Asymptotic properties

We first discuss the asymptotic property of ***β*** estimator given the true ***γ***. As ***β*** is estimated by maximizing the log-likelihood function, we have the following theorem.

##### Theorem 1.

*Assume*
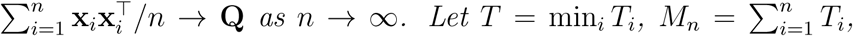 *under the true **γ***, *we have*

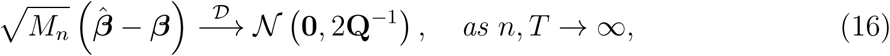

*where*
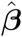 *is the maximum likelihood estimator when the true **γ** is known*.

##### Algorithm 3

A common principal component based method for solving optimization problem (2) under constraint (C2).

**Input:**

**Y**: a list of data where the ith element is a *T_i_* × *p* data matrix

**X**: a *n* × *q* matrix of covariate variables with the first column of ones

**Output:** 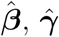

(i) Use Flury (1984) method to estimate Φ and Δ_*i*_ (*i* = 1, …, *n*) for **Y** with *n* groups.
(ii) For *j* = 1, …, *p*, estimate *β* with *γ* = *ϕ_j_*, denoted by *β*^(*j*)^.
(iii) For each *j*, minimize the objective function with

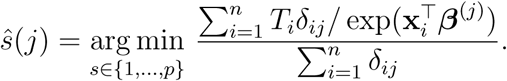
(iv) For those *ŝ*(*j*) = *j*, maximize the variance with

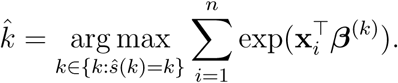
(v) Estimate 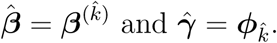

When *p* = 1 and *T_i_* = 1 (for *i* = 1, …, *n*), our proposed model (1) degenerates to a multiplicative heteroscedastic regression model. The asymptotic distribution of
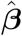 in Theorem 1 is the same as in Harvey (1976). We now establish the asymptotic theory when ***γ*** is estimated from the common principal component approach (Flury, 1984).

##### Theorem 2.

*Assume* Σ_*i*_ = ΓΛ_*i*_Γ^⊤^, *where* Γ = (***γ***_1_, …, **γ**_*p*_) *is an orthogonal matrix and* Λ_*i*_ = diag{λ_*i*1_, …, λ_*ip*_} *with* λ_*ik*_ ≠ λ_*il*_ (*k* ≠ *l*), *for at least one i* ∈ {1, …, *n*}. *There exists k* ∈ {1, …, *p*} *such that for* ∀ *i* ∈ {1, …, *n*},
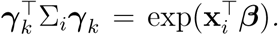 *Let*
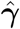 *be the maximum likelihood estimator of **γ**_k_ in Flury (1984). Then assuming that the assumptions in Theorem 1 are satisfied*,
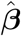 *from Algorithm 3 is*
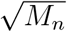-*consistent estimator of **β***.

## 4 Simulation Study

In the simulation study, we generate data from a multivariate normal distribution with *p* = 5 and covariance Σ_*i*_ for sample *i*. We assume the covariance matrices satisfy the common diagonalization assumption, i.e., Σ_*i*_ = ΓΛ_*i*_Γ^*⊤*^, where

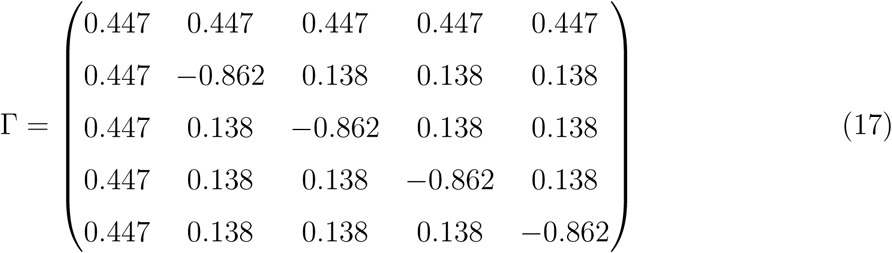

is an orthogonal matrix, and Λ_*i*_ is a diagonal matrix with diagonal elements {λ_*i*1_, …, λ_*ip*_}. For the log-linear model, *X_i_* (*i* = 1, …, *n*) is generated from a Bernoulli distribution with probability 0.5 to be one. Thus *q* = 2 because of the additional intercept column. Two scenarios are tested: (i) the null case with *β*_1_ = 0 and (ii) the alternative case with the second and third eigenvalues satisfying the regression model. For the first cases with *β*_1_ = 0, λ_*ij*_ is generated from a log-normal distribution with mean *β*_0_ and variance 0.5^2^; and for the second case with *β*_1_ ≠ 0, λ_*ij*_ =
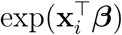 with **x**_*i*_ = (1 *X_i_*)^*⊤*^. The simulation is repeated 200 times.

As demonstrated in Section 3, constraint (C1) yields the component with the lowest normalized data variation. Thus, in this section, we only present the performance under constraint (C2). Under constraint (C2), for higher-order directions, enforcing the orthogonality constraint reduces the parameter search space and increases computation complexity. In this simulation study, we implement both cases with and without the orthogonality constraint. We compare the following methods:

(1) our proposed block coordinate descent method (Algorithms 1 and 2) under constraint (C2) without the orthogonality constraint, denoted as CAP;
(2) our proposed block coordinate descent method (Algorithms 1 and 2) under constraint (C2) with the orthogonality constraint, denoted as CAP-OC;
(3) our method under the complete common principal component model (Algorithm 3) for finding the first *k* projection directions, denoted as CAP-C;
(4) a principal component analysis (PCA) based method, where we apply PCA on each subject and regress each of the first *k* eigenvalues on the covariates, denoted as PCA;
(5) a common principal component method, where we apply common PCA on all subjects using the method in Flury (1984) and regress each of the first *k* eigenvalues on the covariates, denoted as CPCA.

We first evaluate the performance under the null case, i.e., *β*_1_ = 0. *β*_0_’s are set to be ***β***_0_ = (5, 4, 1, −1, −2)^*⊤*^. We present the estimate of *β*’s from CAP and CAP-C over 200 simulations in Figure E.2 in the supplementary material. Our estimate of *β*_1_ is centered around zero with an average of 0.01 under CAP and −0.01 under CAP-C, both much smaller than the corresponding standard errors.

Under the alternative scenario, we set ***β*** as

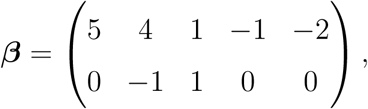

where the first row is for the intercept term (*β*_0_’s). Under this setting, the second and third eigencomponents of the covariance matrices follow the log-linear model (1) and the eigencondition (Condition 1) is satisfied. Table 1 presents the estimate of model coefficients *β*_1_ over 200 simulations with *n* = 100 and *T_i_* = 100. Since the intercept term is not of our study interest, we will not report the results here. For our proposed methods (CAP and CAP-C), the coverage probability is obtained by both the asymptotic variance in Theorem 1 (CP-A) and 500 bootstrap samples (CP-B); while for PCA and CPCA approaches, only CP-B is reported. As the data is generated under the complete common principal component assumption, the CAP-C approach yields the estimate of *β* with the lowest bias. The estimated *β* from CAP (or CAP-OC) for the first direction (the second eigencomponent) is very close to those from CAP-C, and the coverage probability from either the asymptotic variance or bootstrap achieves the designated level (*α* = 0.05). For the second direction (the third eigencomponent), the estimate from CAP has slightly higher bias and the coverage probability is smaller than 0.95. The estimation bias of CAP-OC is higher, due to the orthogonality restriction. The higher bias in the proposed CAP approaches is possibly due to the data manipulation step in Algorithm 2. Both PCA and CPCA do not take into account the covariate information and thus the first two direction estimates are associated with the *β* components corresponding the largest two eigenvalues, even though the first *β* component is zero. The estimate of ***γ*** from CAP, CAP-OC and CAP-C are presented in Table E.1 of Section E in the supplementary material.

**Table 1:** Estimate (Est.) of *β*_1_, as well as standard error (SE), coverage probability with asymptotic variance in Theorem 1 (CP-A) and coverage probability from 500 bootstrap samples (CP-B) from different methods under the alternative hypothesis. All values are computed with *n* = 100 and *T_i_* = 100 over 200 simulations.

| Method | First Direction |  |  | Second Direction |  |  |
| --- | --- | --- | --- | --- | --- | --- |
|  | Est. (SE) | CP-A | CP-B | Est. (SE) | CP-A | CP-B |
| Truth | -1.00 | - | - | 1.00 | - | - |
| CAP | -1.00 (0.03) | 0.950 | 0.950 | 0.81 (0.58) | 0.885 | 0.870 |
| CAP-OC | -1.00 (0.03) | 0.950 | 0.950 | 0.52 (0.84) | 0.730 | 0.715 |
| CAP-C | -1.00 (0.03) | 0.950 | 0.955 | 1.00 (0.03) | 0.975 | 0.960 |
| PCA | -0.02 (0.10) | - | 0 | -0.98 (0.03) | - | 0 |
| CPCA | -0.01 (0.11) | - | 0 | -1.00 (0.03) | - | 0 |

To further assess the finite sample performance of our proposed CAP approach, we vary the number of subjects with *n* = 50, 100, 500, 1000 and the number of observations within subject with *T_i_* = 50, 100, 500, 100. Figure 2 shows the estimate, coverage probability from the asymptotic variance, and the mean squared error (MSE) of model coefficients of the first two directions. From the figure, as both *n* and *T_i_* increase, the estimate of *β*_1_ in both the first and second identified direction converge to the true value. The coverage probability of the first direction is always close to the designated level, and the coverage probability of the second direction converges to 0.95 as both *n* and *T_i_* increase. The MSE of both directions converge to zero. The simulation results demonstrate that our proposed method (CAP) can successfully recover the eigencomponents that possess multiplicative heteroscedasticity. Similar results from CAP-C are shown in Figure E.3 of Section E in the supplementary material.

**Figure 2:**
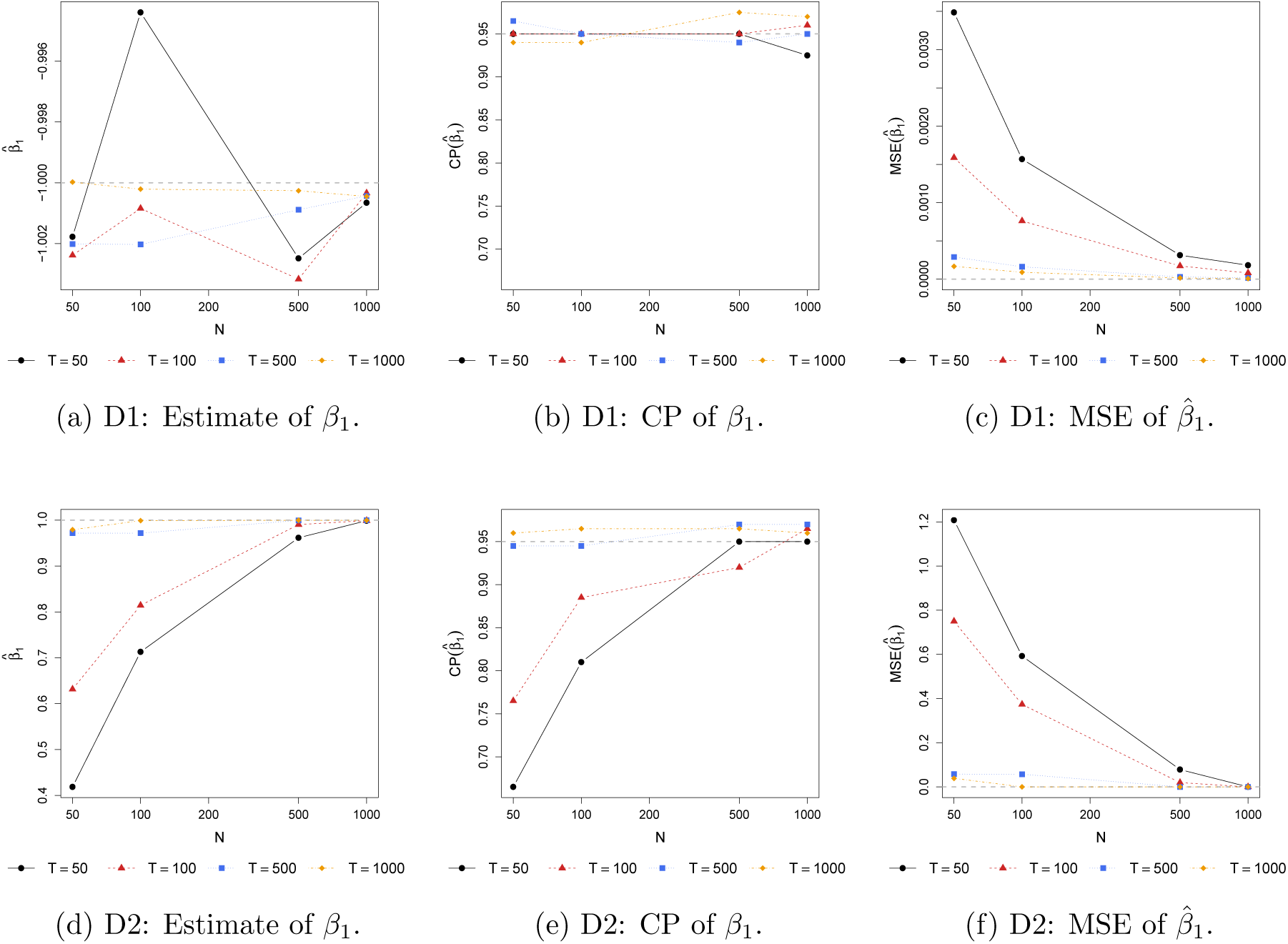
Estimate and coverage probability (CP) with asymptotic variance (Theorem 1) of ***β***_1_ for the first (D1) and second (D2) projection directions, as well as the mean squared error (MSE) of ***β*** estimates under various combination of *n* and *T* values using CAP. The gray dashed lines are the target of estimates in (a) and (d), the designated level 0.95 in (b) and (e), and zero in (c) and (f).

## 5 Resting-state fMRI data example

We apply our proposed method to Human Connectome Project (HCP) resting-state fMRI (rs-fMRI) data. Our dataset includes *n* = 118 healthy young adults (39 aged 22-25 and 79 aged 26-30; 42 female and 76 male) from the most recent S1200 release. The sample size selected here is typical for fMRI studies. The rs-fMRI dataset was preprocessed following the minimal preprocessing pipeline in Glasser et al. (2013). Global signal regression was performed to address whole brain fluctuations typically seen as nuisances (Murphy et al., 2009; Fox et al., 2009). The blood-oxygen-level dependent (BOLD) signals are extracted from *p* = 20 functional brain regions in the default mode network (DMN) (Power et al., 2011) and averaged over voxels within the 5 mm radius. The BOLD time series are temporally correlated, thus we first calculate the effective sample size (ESS) defined by Kass et al. (1998), which infers the equivalent sample size of independent samples,

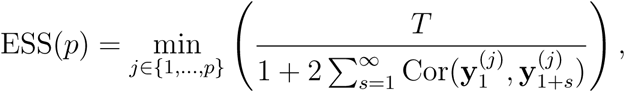

where
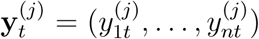 is the data at time *t* of the *j*th brain region from all subjects, for *t* = 1, 2, … and *j* = 1, …, *p*, and *T* = 1200 is the number of time points. We subsample ESS(*p*) = 660 time points (demeaned and variance stabilized (Beckmann and Smith, 2004)) for analysis.

It has been shown that there exists sex discrepancy in functional connectivity in the DMN (Gong et al., 2011; Zhang et al., 2018). In this study, the individual demographic information, i.e., age and sex (both as categorical variables), together with their interaction are considered as the explanatory variables. For age, the category 22-25 is the reference level and labeled as Age1 and 26-30 as Age2; for sex, sex = 1 for male and 0 for female.

Four methods are compared in this study, including *(i)* element-wise correlation regression (Wang et al., 2007); *(ii)* common principal component method, i.e., the CPCA method in Section 4; *(iii)* our CAP-C method; and *(iv)* our CAP method. In the simulation study (Section 4), it is shown that the proposed CAP approach without orthogonal constraint overperforms the approach with orthogonal constraint (CAP-OC). In this real data analysis, we employ the CAP approach and include a *post hoc* procedure to examine the orthogonality among the identified projection directions.

For the element-wise correlation regression, each off-diagonal element in the correlation matrix is Fisher *z*-transformed and multiple testing adjustment is performed following the Benjamini and Hochberg (1995) procedure to control the false discovery rate (FDR). None of the FDR corrected p-values are significant at level 0.05. See Figure F.1 and Figure F.2 in the supplementary material for the raw and FDR corrected *p*-values.

We present the estimated regression coefficients (together with 95% confidence intervals) of first ten common PCs from the CPCA approach in Figure F.3 in the supplementary material. From the figure, only the model coefficients of the fourth component (CPC4) are significant, indicating that not all of the top PCs are related to either age or sex. Under the same common PCA assumption, CAP-C directly discovers the PCs that are relevant to the covariates. Thus, the first component identified by CAP-C is CPC4. Figure F.4 shows the estimated model coefficients (and 95% confidence intervals from 500 bootstrap samples) of the top seven discovered PCs.

Our CAP approach discovers five projection directions, where the number five is chosen based on the average DfD (see Figure F.5 in Section F of the supplementary material), and the orthogonality of these five directions are verified in Figure F.10. Figure 3 exhibits the model coefficients (and 95% confidence intervals from 500 bootstrap samples) of the five projection directions. From the figures, for each identified projection direction, at least one of the covariates is significant. We use D1 as an example, which presents a significant age effect. To interpret the loadings, Figure 4a shows the loading profile, and six brain regions have the loading magnitude greater than 0.2 (see Figure 4b in the brain map). This suggests that the connectivity between these brain regions show significant difference in the comparison (see Figure F.9 in the supplementary material for the scatter plot). In in the supplementary material, additional loading plots and brain maps are available in Figures F.7 and F.8, comparisons under different contrasts are shown in Figure F.6. To compare with CAP-C, Table F.1 in the supplementary material displays the similarity (similarity between −1 and 1, and 0 indicates orthogonal) of the projection directions to the common PCs from CAP-C. The two significant PCs identified by CAP-C, CPC4 and CPC18, have similarity greater than 0.6 to the projections D2 and D4 identified by CAP, respectively.

**Figure 3:**
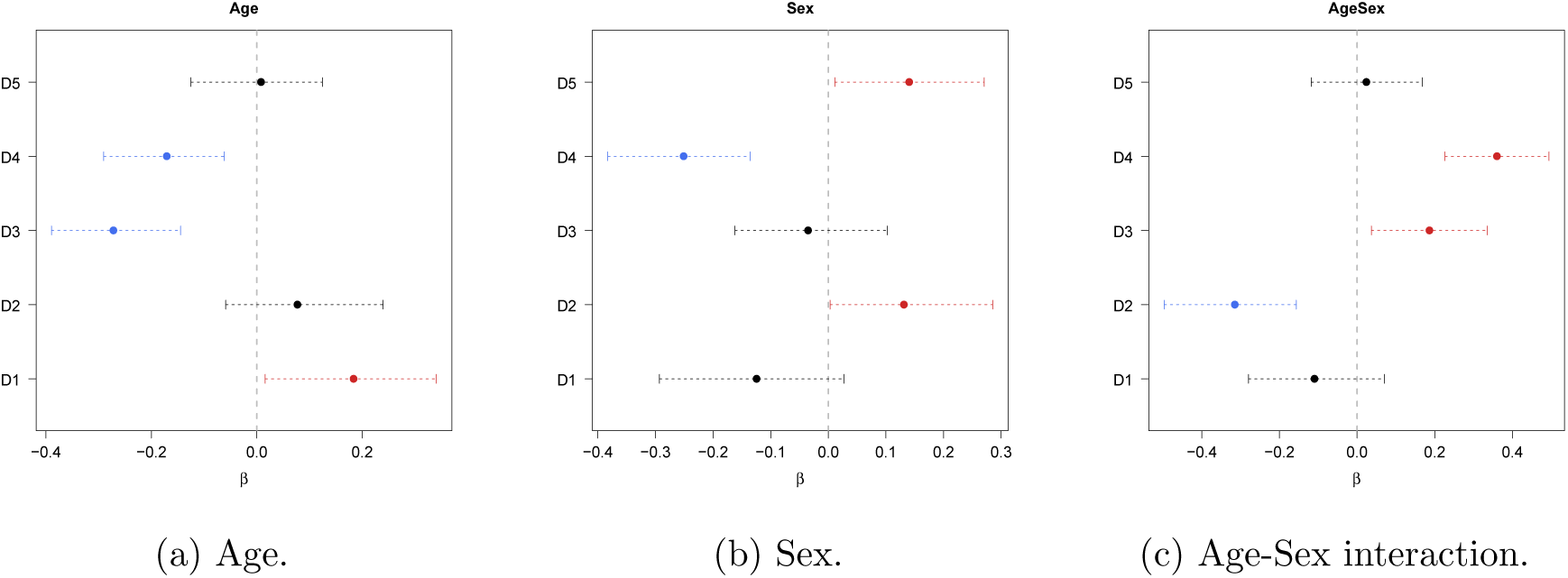
Estimated model coefficients and 95% bootstrap confidence intervals of the six identified projection directions by CAP.

**Figure 4:**
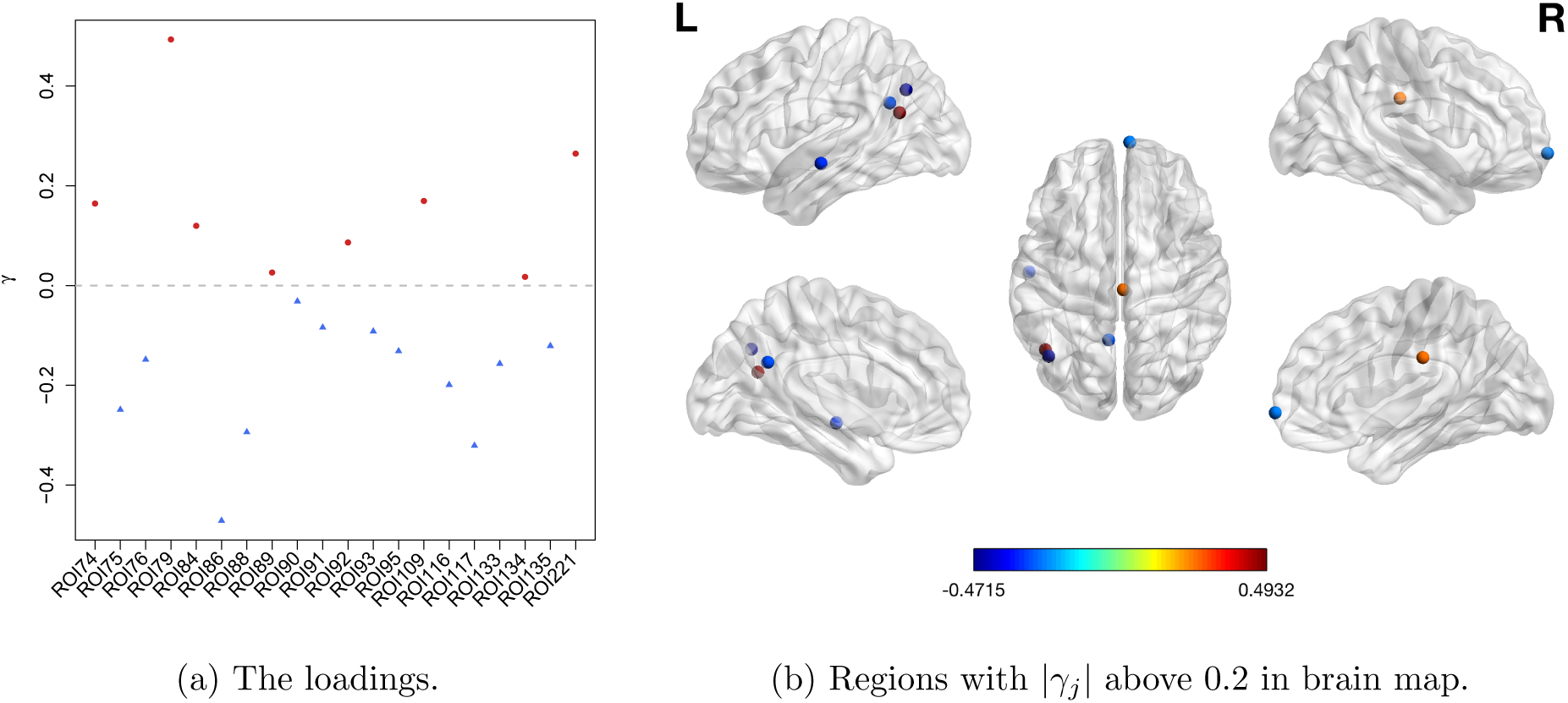
The loading profile and brain regions with absolute loading greater than 0.2 in projection direction D1 identified by CAP.

To study the reliability of our proposed methods, we apply the estimated projections to three other scanning sessions of resting-state fMRI data acquired from the same subjects. Figure F.11 in the supplementary material shows the estimated model coefficients and 95% bootstrap confidence intervals and Figure F.12 presents the comparisons under different contrasts. From the figures, the estimate and significance are very similar to the result presented in Figure 3, which validates the existence of difference between age groups and/or sex within these five components (also known as brain subnetworks) of the DMN.

## 6 Discussion

In this study, we introduce a Covariate Assisted Principal regression model for multiple co-variance matrix outcomes. Our approach allows the identification of projection directions that are associated with the explanatory variables or covariates. Under certain regularity conditions, our proposed estimators are asymptotically consistent. Using extensive simulation studies, our model shows high estimation accuracy. Applied to resting-state fMRI studies, our method avoids the massive number of hypothesis testing suffered in the element-wise regression approach.

One challenge in modeling covariance matrices directly is having a constraint of positive definiteness. Via projections, the study of a positive definite matrix is decomposed into modeling the eigenvalues in orthogonal spaces. This relaxes the constraint and preserves geometric interpretation. The existing spectral decomposition based methods rely on the assumption that there exists a common diagonalization of the covariance matrices. In practice, this can be unrealistic, especially when *p* is large. Researchers are often more interested in studying a subset of the components related to the covariates. Though CAP enables identification of a small set of components, the theoretical analysis is challenging without the complete common diagonalization regularity condition in CPCA. One future direction will consider relaxing these assumptions.

The current framework assumes the dimension of the data, *p*, is fixed and less than both the number of observations within a subject and the number of subjects. Another future direction is to extend the method to settings of large *p*, small *n*.

## Supplementary Materials

### A Theory and Proof

#### A.1 A proposition for Algorithm 1 and proof of Proposition 1

##### Proposition A.1.

*Suppose the vector* **x** *∈* ℝ^*p*^ *is subject to the restriction*

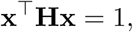

*where matrix* **H** *is positive definite. Then*, *the stationary points and values of* **x**^⊤^ **Ax** *are the eigenvectors and values of* **A** *with respect to* **H**.

##### Proof.

The Lagrangian of the optimization problem is

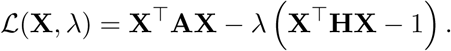

Taking partial derivatives gives

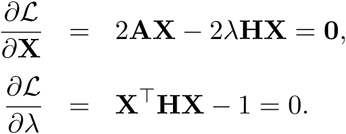

Then

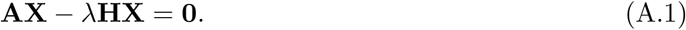

Thus the solution (**X**, λ) is the eigenvector and eigenvalue of **A** with respect to **H**. The proof of Proposition 1 is straight forward by replacing **H** with **I**.

To find the eigenvectors and eigenvalues of **A** with respect to **H**, we first assume **x**_0_ is a solution eigenvector that has Euclidean norm 1, i.e,
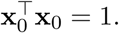
Since **H** is positive definite, let **x** = **H**^−1/2^**x**_0_, then

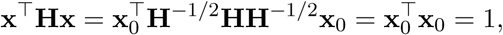

which satisfies the constraint condition. Replace **X** with **x** = **H**^−1/2^**x**_0_ in (A.1),

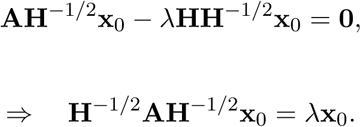

Therefore, **x**_0_ is the eigenvector of matrix **H**^*−*1*/*2^ **AH**^*−*1*/*2^.

In Algorithm 1, for the (*s* + 1)the step,

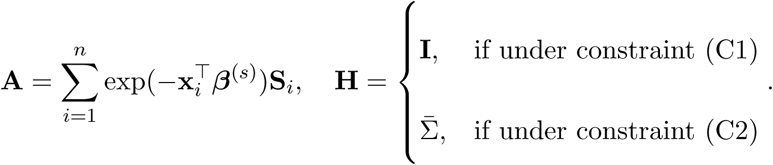

We can first find the eigenvectors of **H**^*−*1*/*2^ **AH**^*−*1*/*2^, left multiplied by **H**^*−*1*/*2^, solve for ***β*** using formula (5). The update of *γ* and ***β*** will be the pair that jointly minimizes the objective function.

#### A.2 Details of Algorithm 2

In Algorithm 2, with the new data
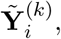
we need to solve the following optimization problem

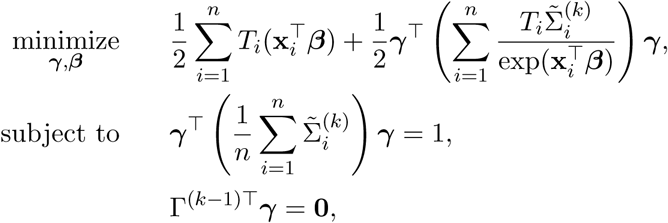

where
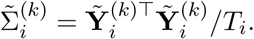
With given (or an initial) ***β***, let

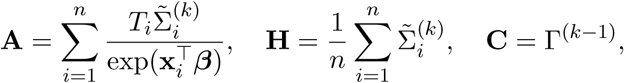

we first apply the solution in Rao (1964, 1973) to find the stationary points, which are the eigenvectors of (**I** − **P**)**A** with respect to **H**, where **P** = **C**(**C**^⊤^ **H**^−1^ **C**)^−^ **C**^⊤^ **H**^−1^ is the projection operator onto 𝓜(**C**) (the linear manifold spanned by **C**). For each eigenvector, find the solution for ***β*** using the formula in Algorithm 1. The update of ***γ*** and ***β*** will be the pair that jointly minimizes the objective function.

### B Proof of Theorem 1

#### Proof.

With true ***γ***, our proposed estimator of ***β*** is the maximum likelihood estimator (MLE). Therefore, the asymptotic results of MLE can be applied.

For subject *i* (*i* = 1, …, *n*) observation *t* (*t* = 1, …, *T_i_*), the log-likelihood function (with a constant difference) is

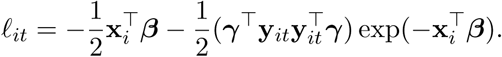

For the full dataset, let *M_n_* = 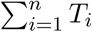 and

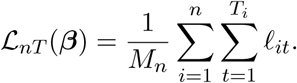

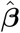 is the solution to 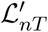 = 0. We expand the function at the true parameter ***β***_0_ as

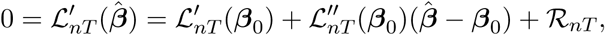

where 𝓡_*nT*_ is the residual term.

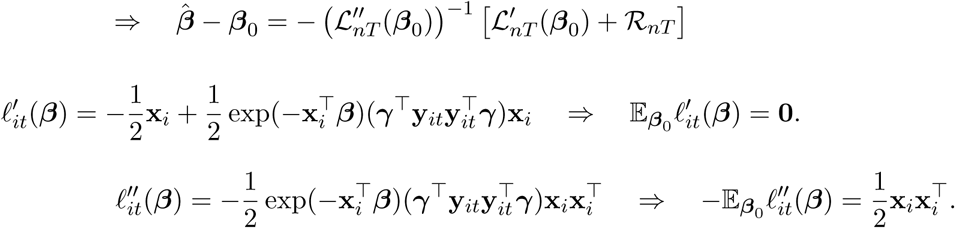

Under the assumption that 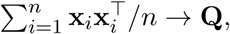

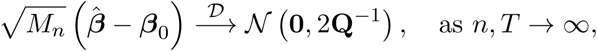

where *T* = min*_i_ T_i_*.

### C Proof of Theorem 2

#### Proof.

We propose to estimate ***γ*** and ***β*** by maximizing the likelihood function. Under the complete common principal component assumption, Flury (1986) showed the asymptotic distribution of 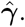 Together with the conclusion of Theorem 1, the consistency of 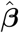 follows.

### D Toy examples

We use three examples to demonstrate the property of the two considered constraints. Assume *X_i_* is generated from a Bernoulli distribution with probability 0.5 to be 1.

#### D.1 Example I

Let ***β***_1_ = (2, 3)*⊤* and ***β***_2_ = (2,−3)*⊤*, and assume Σ_*i*_ = ΓΛ*_i_*Γ^⊤^, where

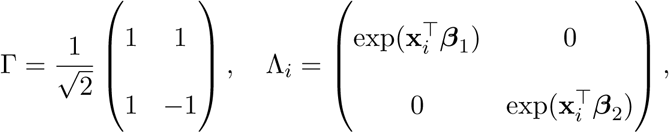

where **x**_*i*_ = (1, *X_i_*)*⊤*. When *X_i_* = 1, Σ _*i*_ =
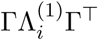 with
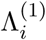 = diag{exp(5), exp(*−*1)}, and when *X_i_* = 0, Σ_*i*_ =
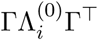 with
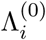 diag{exp(2), exp(2)}, where the projection onto the first eigenspace contains larger variation in the data. We generate **y***_it_*’s from the multivariate normal distribution with mean zero and covariance matrix Σ*_i_*, for *t* = 1, …, *T_i_* = 100 and *i* = 1, …, *n* = 100.

Then Γ^⊤^**y***_it_* follows the multivariate normal distribution with covariance matrix Λ*_i_*. Figures D.1a presents the contour plot of the objective function in 2 with ***β***^∗^ = ***β***_1_. Under (C1), from Algorithm 1, the solution for ***γ*** is the eigenvector corresponding to the minimum eigenvalue of matrix
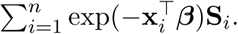 Constraint (C2) regulates the shape of the constraint set by the average sample covariance matrix.

#### D.2 Example II

Let ***β***_1_ = (1, 0)^⊤^ and ***β***_2_ = (−1, 0)^⊤^, which is the null scenario of ***β***. The rest parameter settings are the same as in Example I. Under this scenario, exp
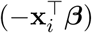is a constant, and thus the constraint set under (C2) is parallel to the contour plot of the objective function under the true ***β*** (see Figure D.1b). Therefore, the estimate of ***γ*** can be any value in the constraint set.

#### D3 Example III

Let ***β*** = (1, −3)^⊤^, and Σ_*i*_ = ΓΛ*_i_*Γ^⊤^, where Λ*_i_* =
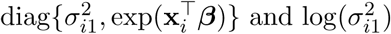 follows a normal distribution with mean two and standard deviation one. The rest parameters are set to be the same as in Example I. In this example, the component with lower variation is relevant to the covariate *X*. Figure D.1c shows the contour plot under the true ***β***. Both constraints identify the second component as the estimator of ***γ***.

Using Examples I to III, we conclude that under constraint (C1), the proposed method yields the estimate of the component with the lowest variation in the data; while constraint (C2) identifies the component that satisfies the model assumption (1).

**Figure D.1:**
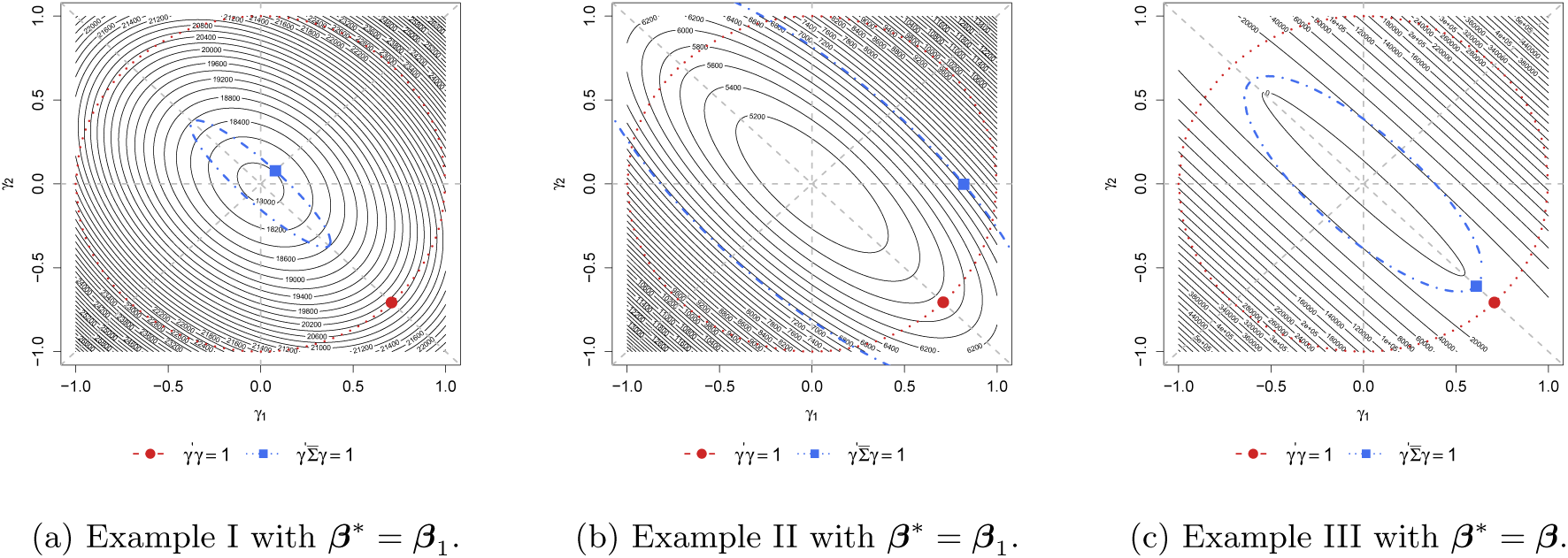
The contour plot of the negative log-likelihood function in (a) Example I, (b) Example II and (c) Example III. The blue curve and point are the constraint (C2) function and the estimate, respectively; and the red are under constraint (C1).

### E Additional Simulation Results

We first use a simulated example to demonstrate the performance of the “deviation from diagonality” metric defined in (8). The data is generated following the alternative scenario in Section 4. Figure E.1a shows the average DfD and Figures E.1b and E.1c are the boxplot of individual DfD, where the *γ*’s are estimated using our proposed CAP method. From the figures, for all samples, when moving to the third component, the DfD value jumps to over 10^6^. Thus, two is the proper number of components to be chosen, which is the same as the truth. Therefore, the proposed average DfD is an appropriate metric to chose the number of projection directions.

Under the null case, we present the estimate of *β*’s from CAP and CAP-C over 200 simulations in Figure E.2. As demonstrated in the toy example II in Section D, under constraint (C2), our method could not identify the principal direction of projection, and thus the estimate of *β*_0_ from CAP and CAP-C varies according to the estimated *γ*. However, the estimate of *β*_1_ is centered around zero with an average of 0.01 (SE: 0.20) under CAP and −0.01 (SE: 0.15) under CAP-C.

**Figure E.1:**
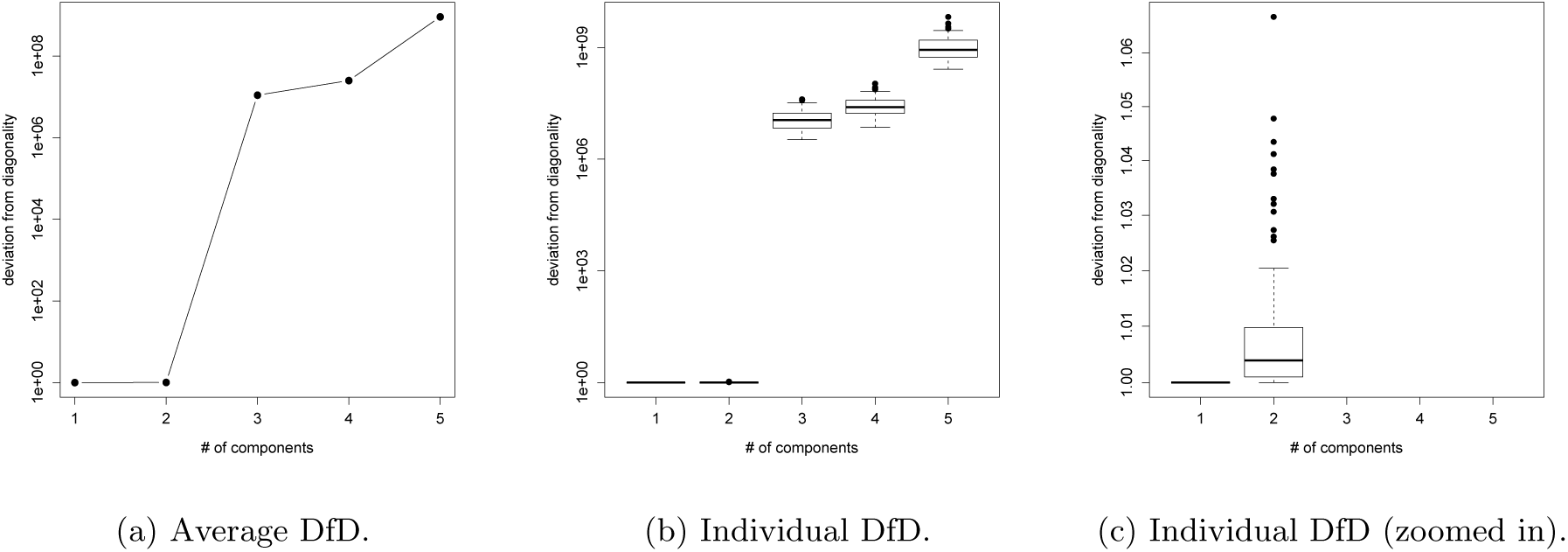
Average and individual “deviation from diagonality” of a simulated example.

**Figure E.2:**
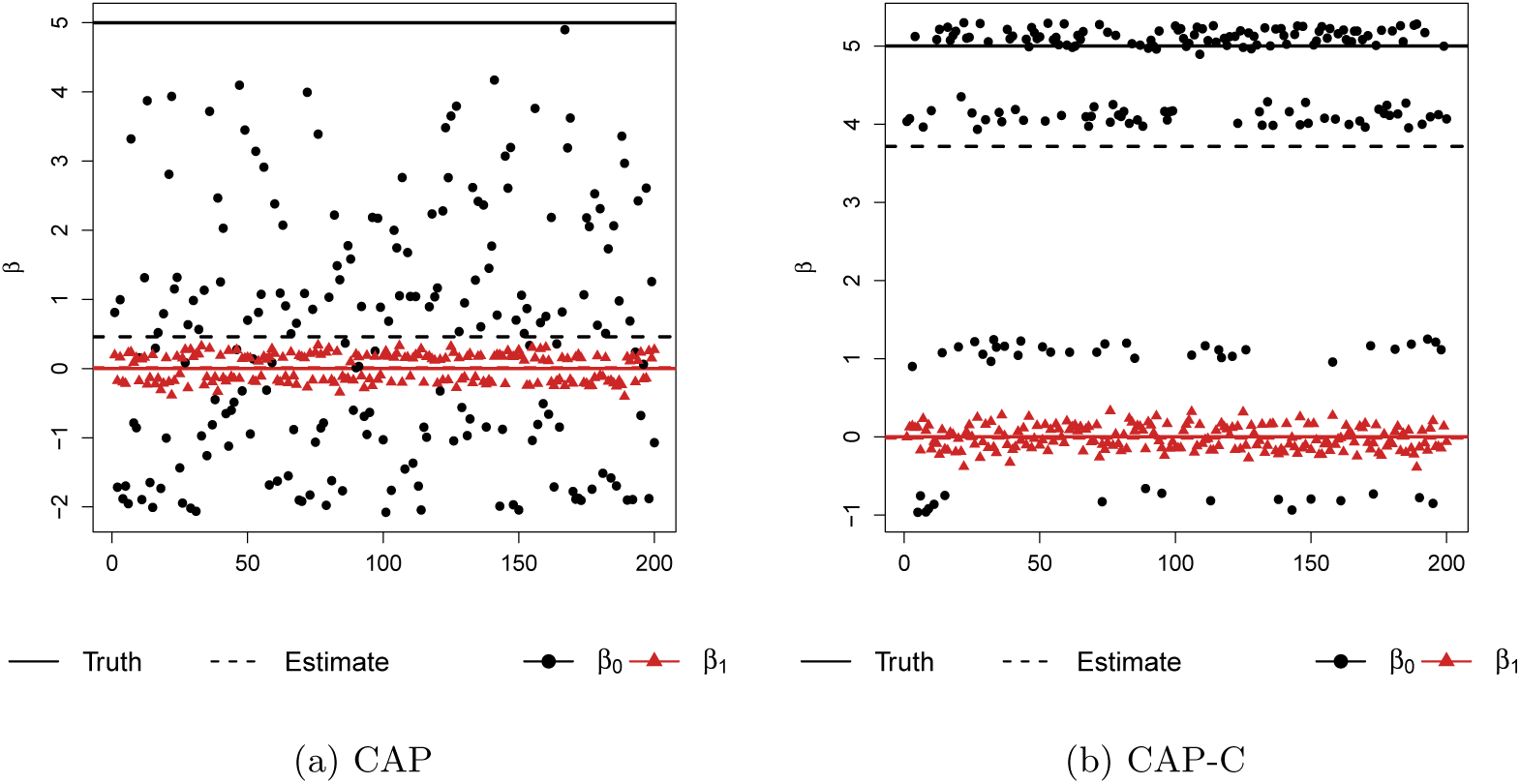
Estimate of *β*_0_ and *β*_1_ in the 200 simulations from (a) CAP and (b) CAP-C methods with *n* = 100 and *T_i_* = 100 under the null case.

Table E.1 presents the estimate of *γ* using CAP and CAP-C methods with *n* = 100 and *T_i_* = 100. Both methods yield correct identification of the two principal directions. CAP-C attains lower bias and variation, which is optimal under the complete common principal component assumption.

**Table E.1:** Estimate (standard error) of *γ* under the alternative scenario with *n* = 100 and *T_i_* = 100.

| | $\gamma$ | Truth | CAP | CAP-OC | CAP-C |
| --- | --- | --- | --- | --- | --- |
| First Direction | $\gamma_1$ | 0.45 | 0.42 (0.125) | 0.42 (0.125) | 0.45 (0.002) |
| | $\gamma_2$ | -0.86 | -0.82 (0.057) | -0.82 (0.057) | -0.86 (0.002) |
| | $\gamma_3$ | 0.14 | 0.13 (0.050) | 0.13 (0.050) | 0.14 (0.004) |
| | $\gamma_4$ | 0.14 | 0.13 (0.161) | 0.13 (0.161) | 0.14 (0.002) |
| | $\gamma_5$ | 0.14 | 0.14 (0.210) | 0.14 (0.210) | 0.14 (0.002) |
| Second Direction | $\gamma_1$ | 0.45 | 0.43 (0.085) | 0.43 (0.111) | 0.45 (0.002) |
| | $\gamma_2$ | 0.14 | 0.15 (0.110) | 0.13 (0.157) | 0.14 (0.003) |
| | $\gamma_3$ | -0.86 | -0.76 (0.282) | -0.61 (0.423) | -0.86 (0.001) |
| | $\gamma_4$ | 0.14 | 0.13 (0.087) | 0.11 (0.171) | 0.14 (0.003) |
| | $\gamma_5$ | 0.14 | 0.06 (0.288) | -0.06 (0.408) | 0.14 (0.002) |

Figure E.3 shows the estimate of *β* using CAP-C as both *n* and *T_i_* increases. As CAP-C correctly identifies the two components that satisfy the model assumption (1), the estimate of *β* is close to the true value and the coverage probability reaches the designated level under all combinations of *n* and *T_i_* values.

**Figure E.3:**
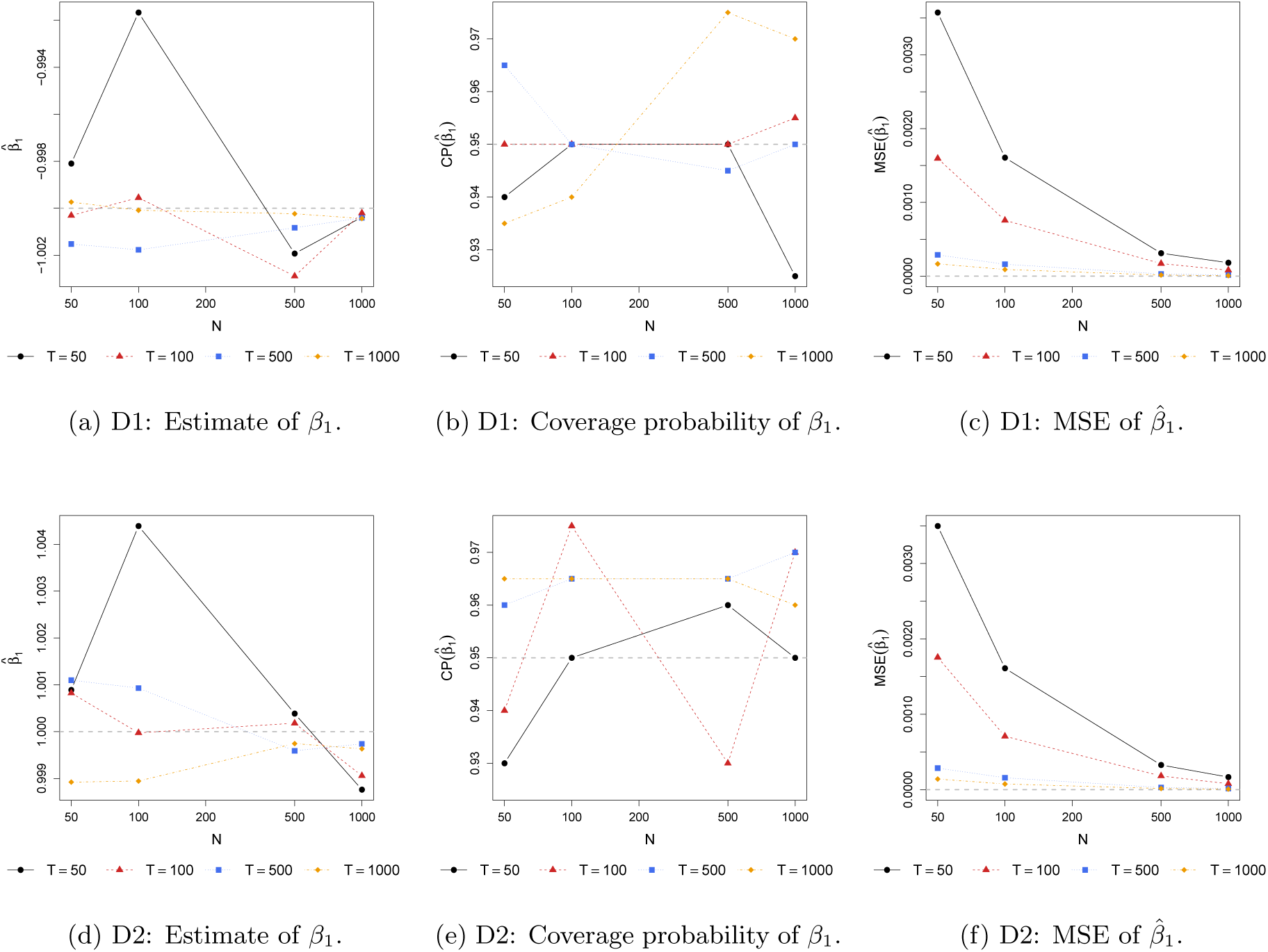
Estimate and coverage probability (CP) with asymptotic variance (Theorem 1) of *β*_1_ for the first (D1) and second (D2) projecting direction, as well as the mean squared error (MSE) of *β* estimates under various combination of *n* and *T* values using CAP-C. The gray dashed line in (a) and (d) are the target of estimates and zero in the rest.

**Figure F.1:**
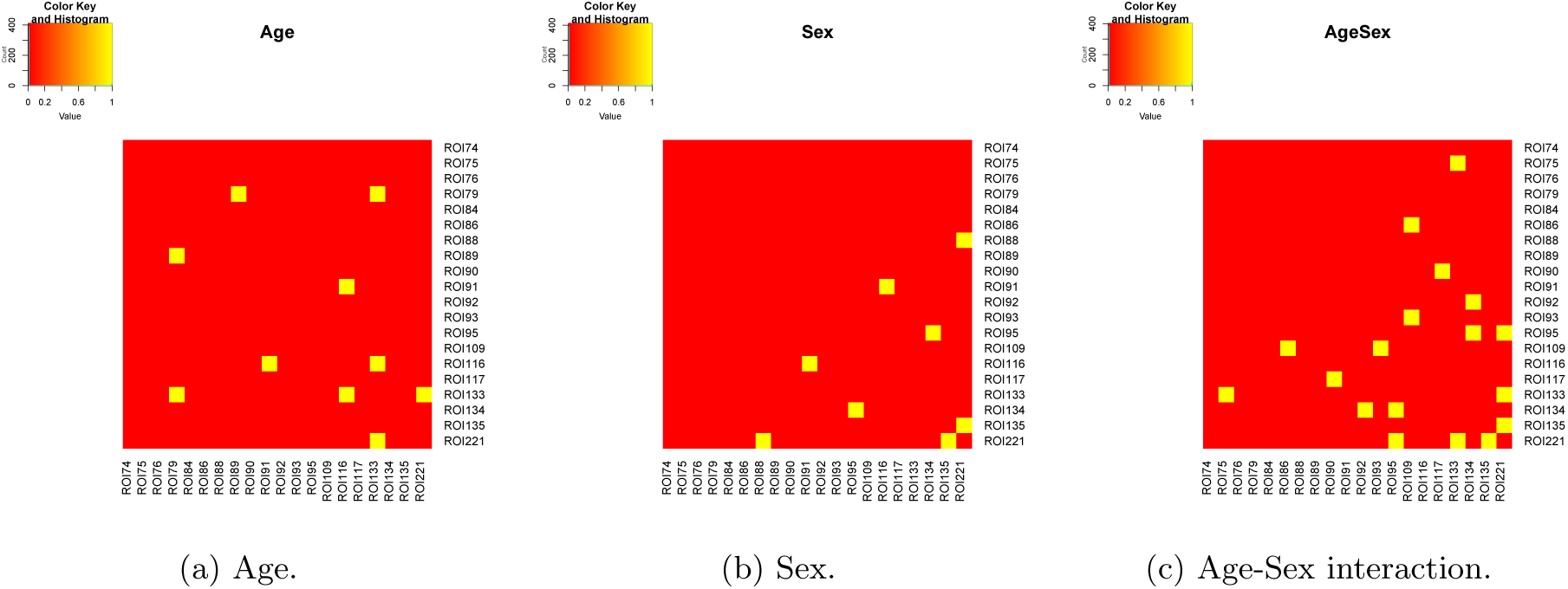
Significance of model coefficients with original *p*-value at level of 0.05 in the element-wise correlation regression. The yellow elements are significant, and the red are not.

### F Additional Real Data Analysis Results

#### F.1 The element-wise regression approach

Figure F.1 shows the significance of model coefficients with original *p*-value less than 0.05 in the element-wise regression analysis. Figure F.2 shows the significance after multiple testing correction, where all become insignificant.

#### F.2 The CPCA approach

We present the estimated model coefficients (together with 95% confidence interval from the regression model) of the first ten common PCs from the CPCA approach in Figure F.3. From the figure, the model coefficients of CPC5, CPC6 and CPC7 are not significant, indicating that brain connectivity within the corresponding brain network does not show any difference when comparing age and sex groups. The CAP-C method builds on the common diagonalization assumption as in the CPCA approach, but targets on the PCs that satisfy the log-linear model assumption.

**Figure F.2:**
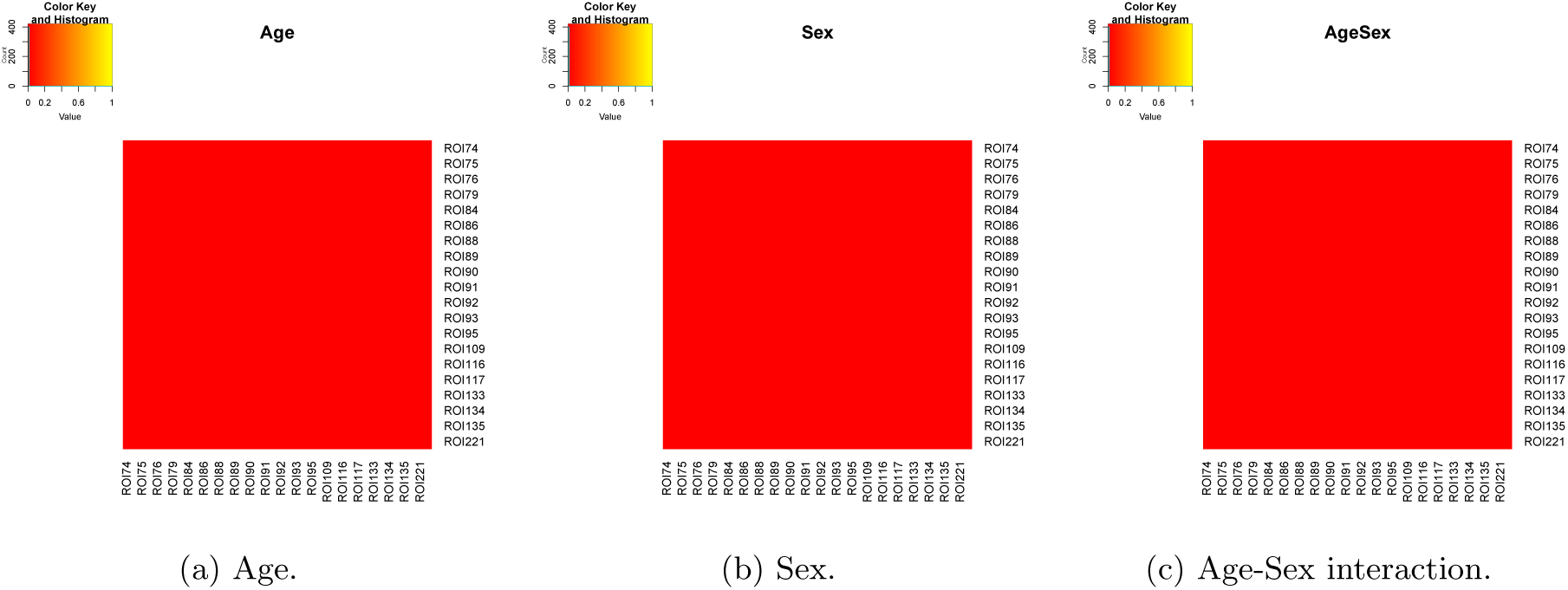
Significance of model coefficients with adjusted *p*-value at level of 0.05 in the element-wise correlation regression. The yellow elements are significant, and the red are not.

**Figure F.3:**
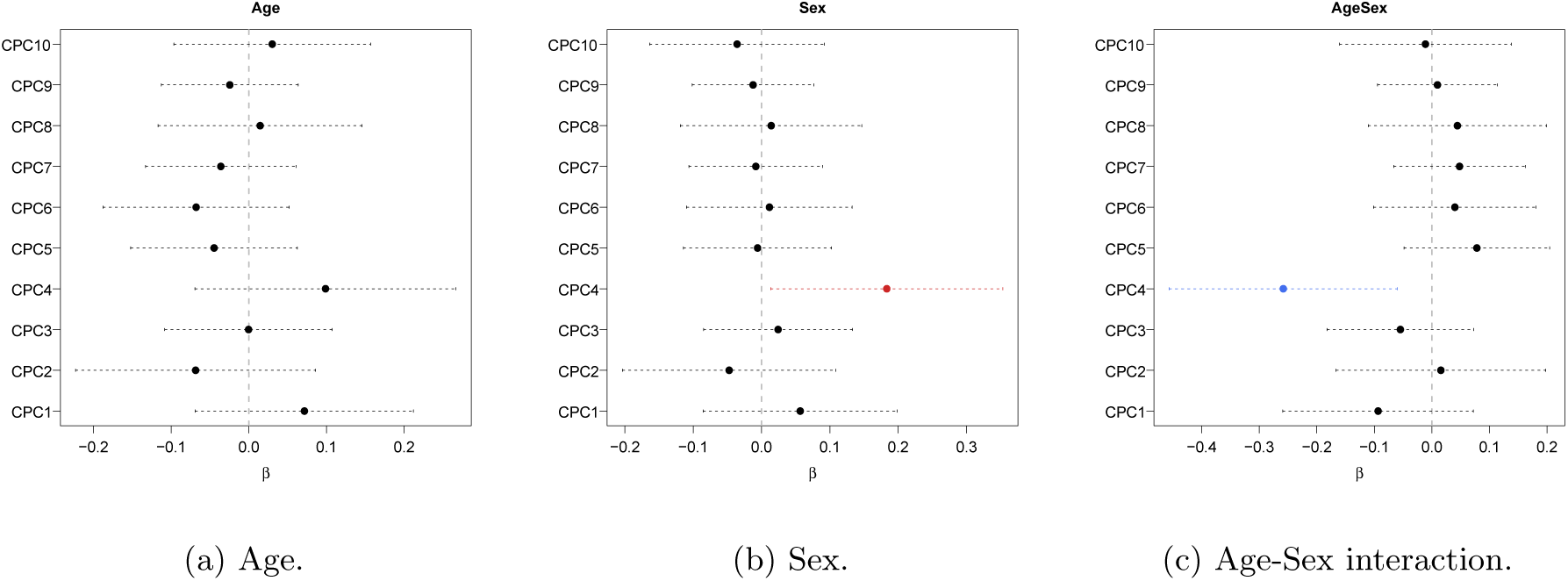
Estimated model coefficients and 95% confidence interval of the first ten common PCs in the CPCA approach.

**Figure F.4:**
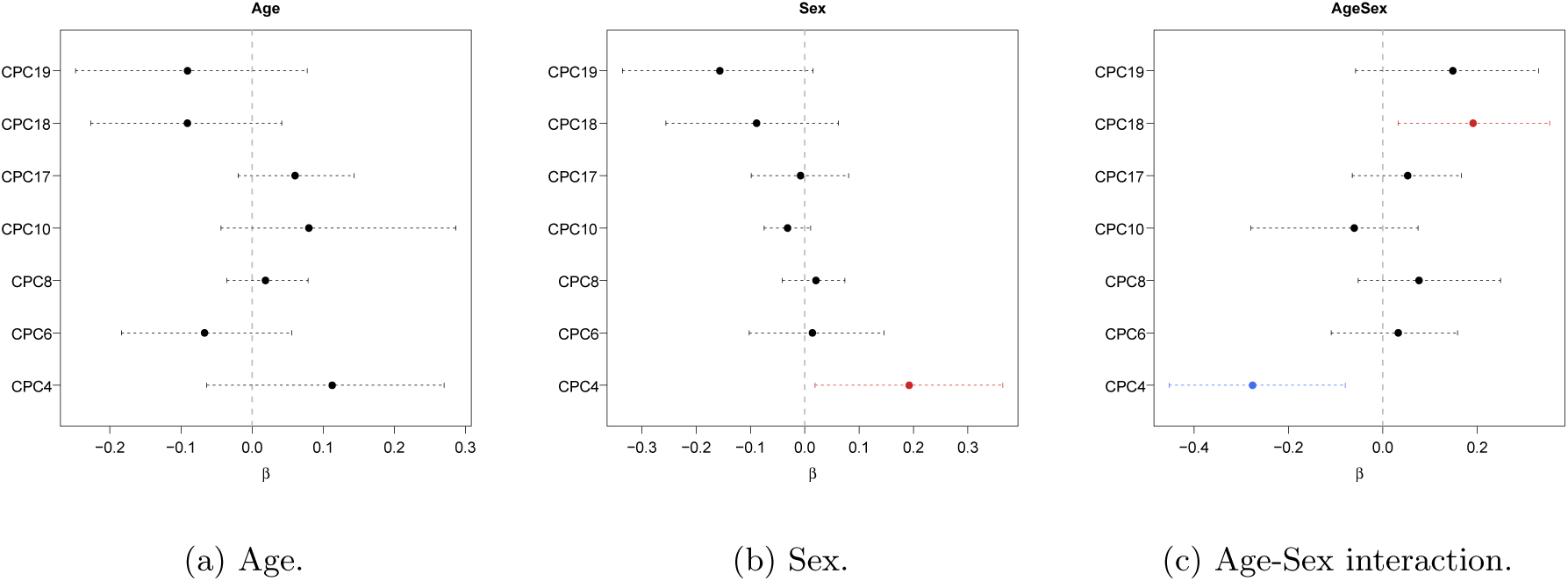
Estimated model coefficients and 95% bootstrap confidence interval of the three identified common PCs from the CAP-C approach.

#### F.3 The CAP-C approach

Figure F.4 shows the estimated model coefficients (and 95% confidence interval from 500 bootstrap samples) of the three discovered PCs, which also satisfy the eigenvalue condition (Condition 1). Though CPC3 has significant coefficient in sex, the corresponding eigenvalue condition is violated and thus is not identified by the CAP-C approach.

#### F.4 The CAP approach

Figure F.5 presents the average and individual “deviation from diagonality” of the first seven projection directions in the real data analysis. We observe a sudden jump on the sixth direction, therefore we choose the first five components.

Figure F.7 presents the loadings of the five projection directions from the CAP approach, and Figure F.8 is the visualization of the loadings in the brain map. Figure F.9 shows the scatter plot of the model outcome log
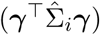by age and sex group for the five projection directions from the CAP approach. From the figure, we observe the interaction effect in D2, D4 and D5.

**Figure F.5:**
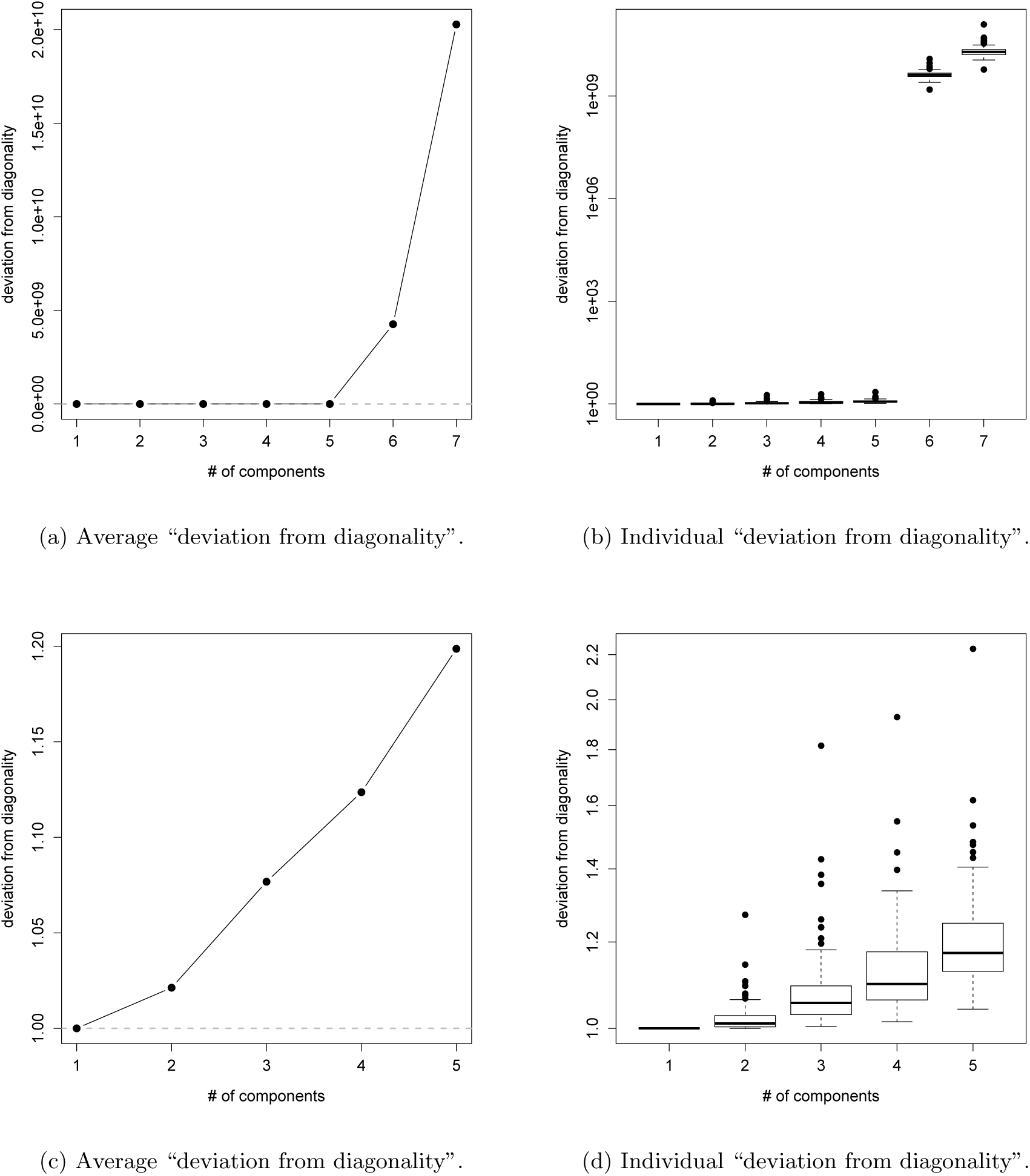
The average and individual “deviation from diagonality” of the first seven ((a)-(b)) and first five ((c)-(d)) projection directions in the real data analysis.

**Figure F.6:**
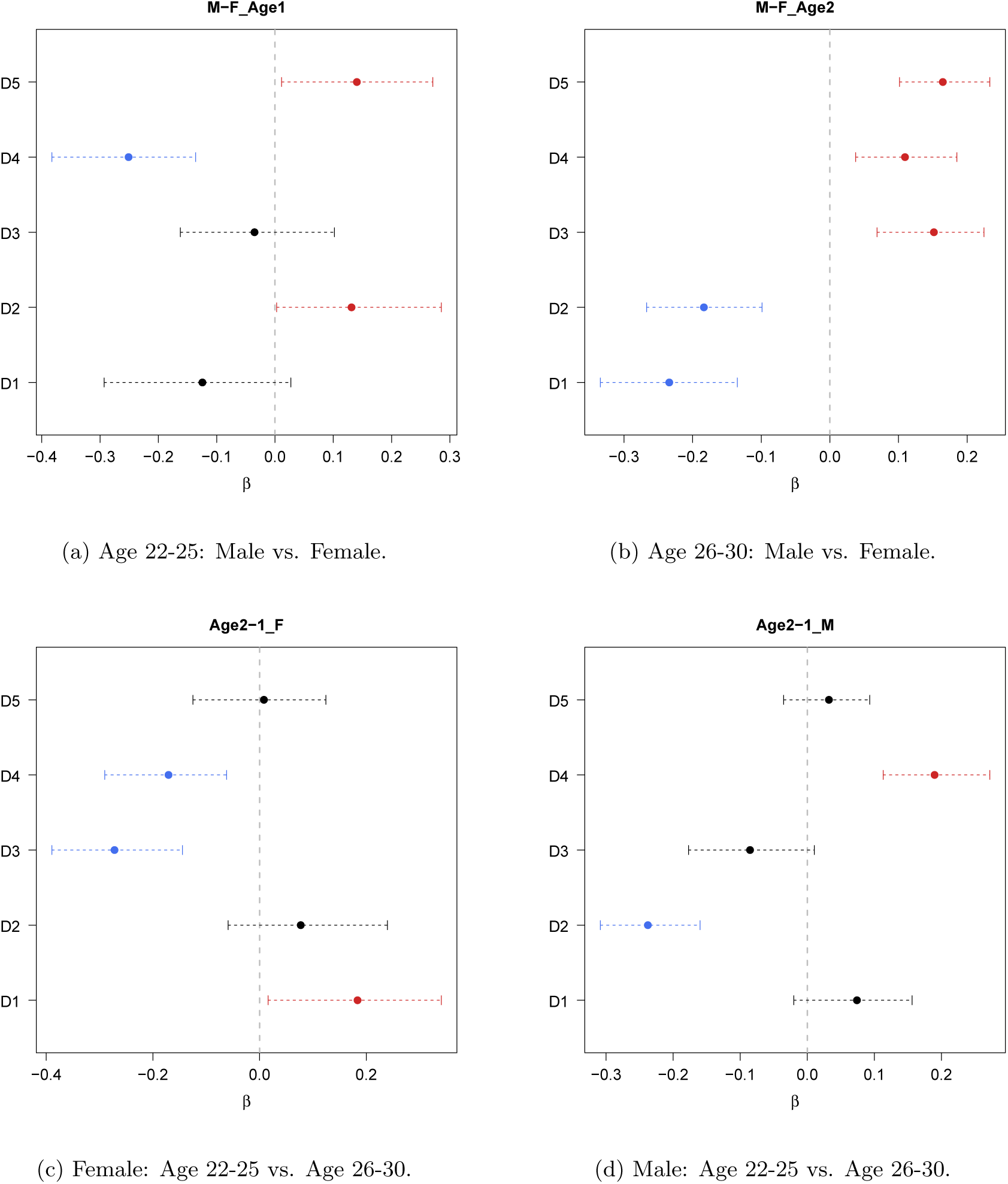
Pair-wise comparison of the five identified projection directions from the CAP approach. The confidence interval is obtained from 500 bootstrap sample.

Table F.1 displays the similarity (similarity between −1 and 1, and 0 indicates orthogonal) of the projecting directions to the PCs from CAP-C. The proposed CAP approach recovers the three PCs with high similarity and detects three additional. Using the definition in Krzanowski (1979), the similarity between the two spaces discovered by CAP and CAP-C is 0.386, indicating that the space spanned by the seven identified PCs from CAP-C is different from the one spanned by the five components discovered by CAP.

**Figure F.7:**
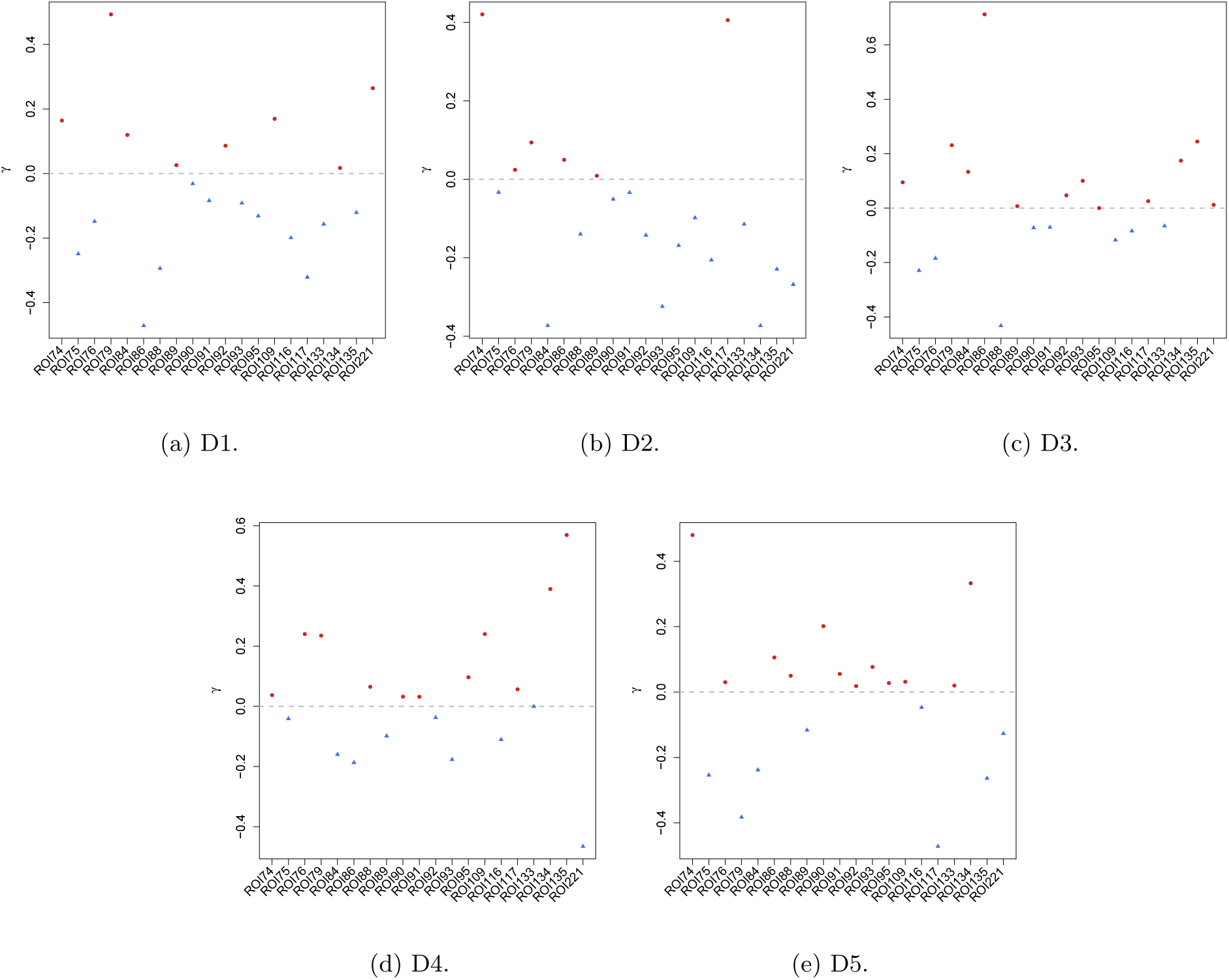
The loadings of the five projection directions from the CAP approach.

**Figure F.8:**
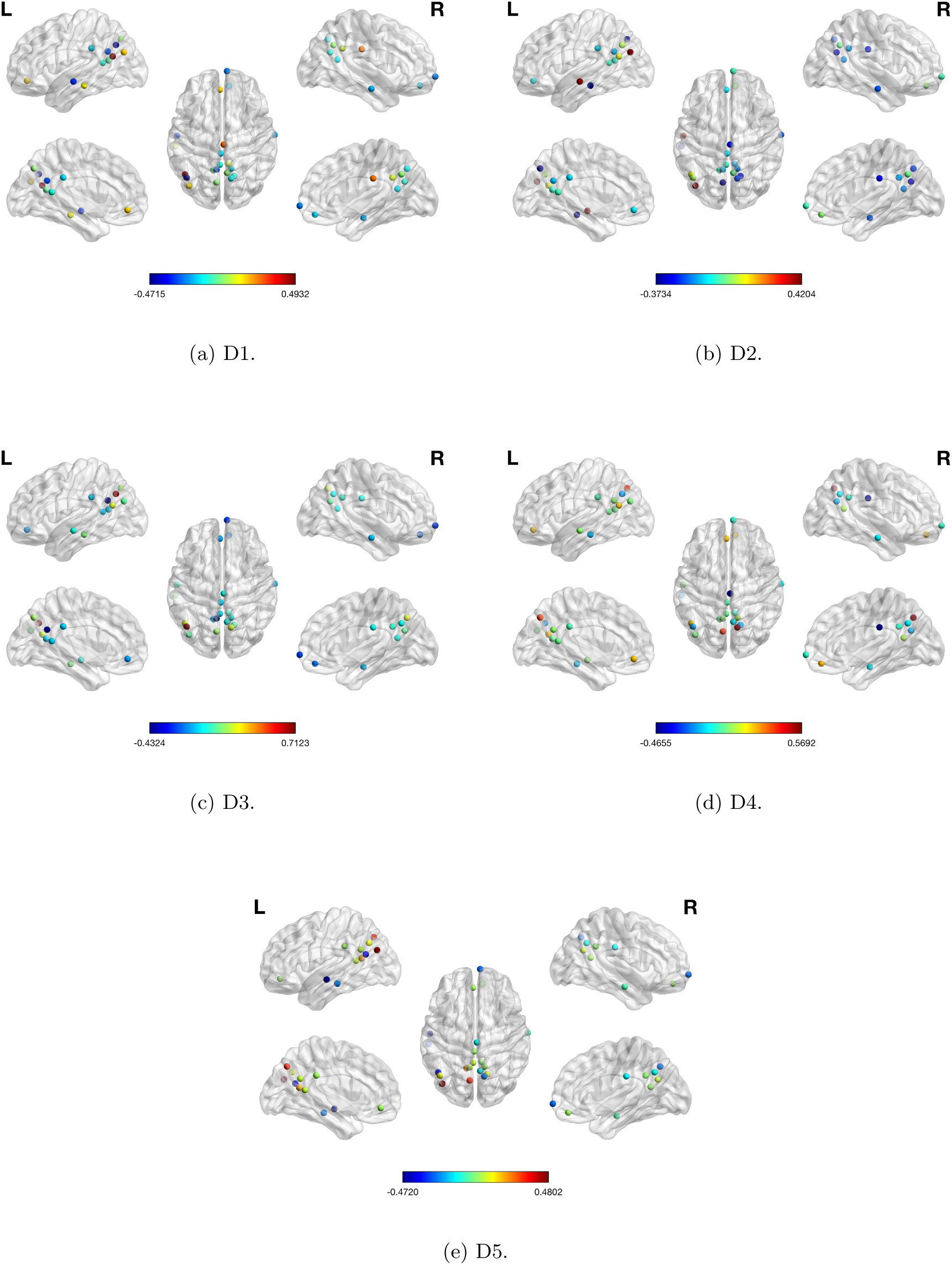
The loading map of the five projection directions from the CAP approach. The color legend indicates the value of *γ*.

**Figure F.9:**
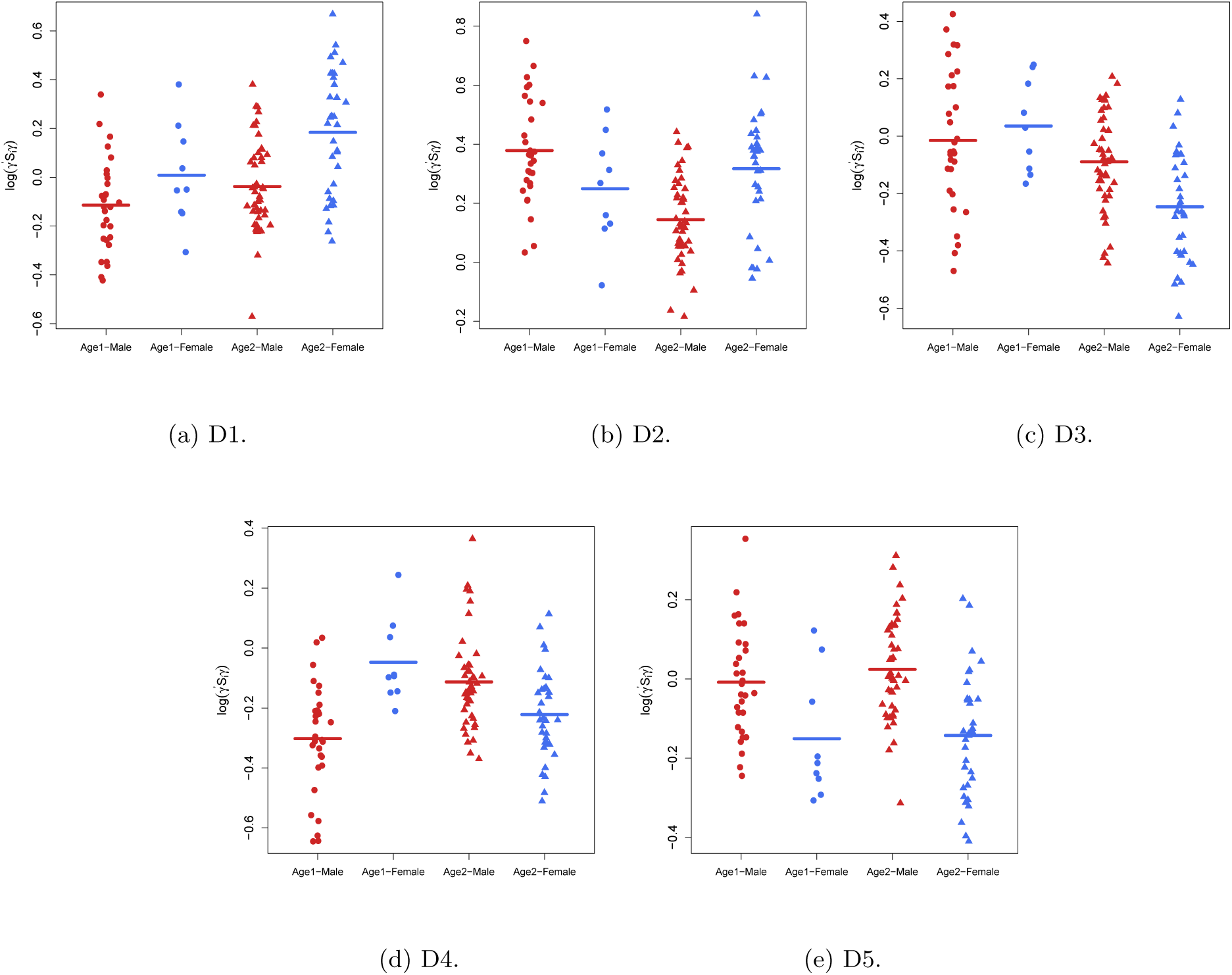
Scatter plot of the outcome in the log-linear model (log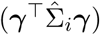) by age and sex groups for the five projection directions from the CAP approach.

**Figure F.10:**
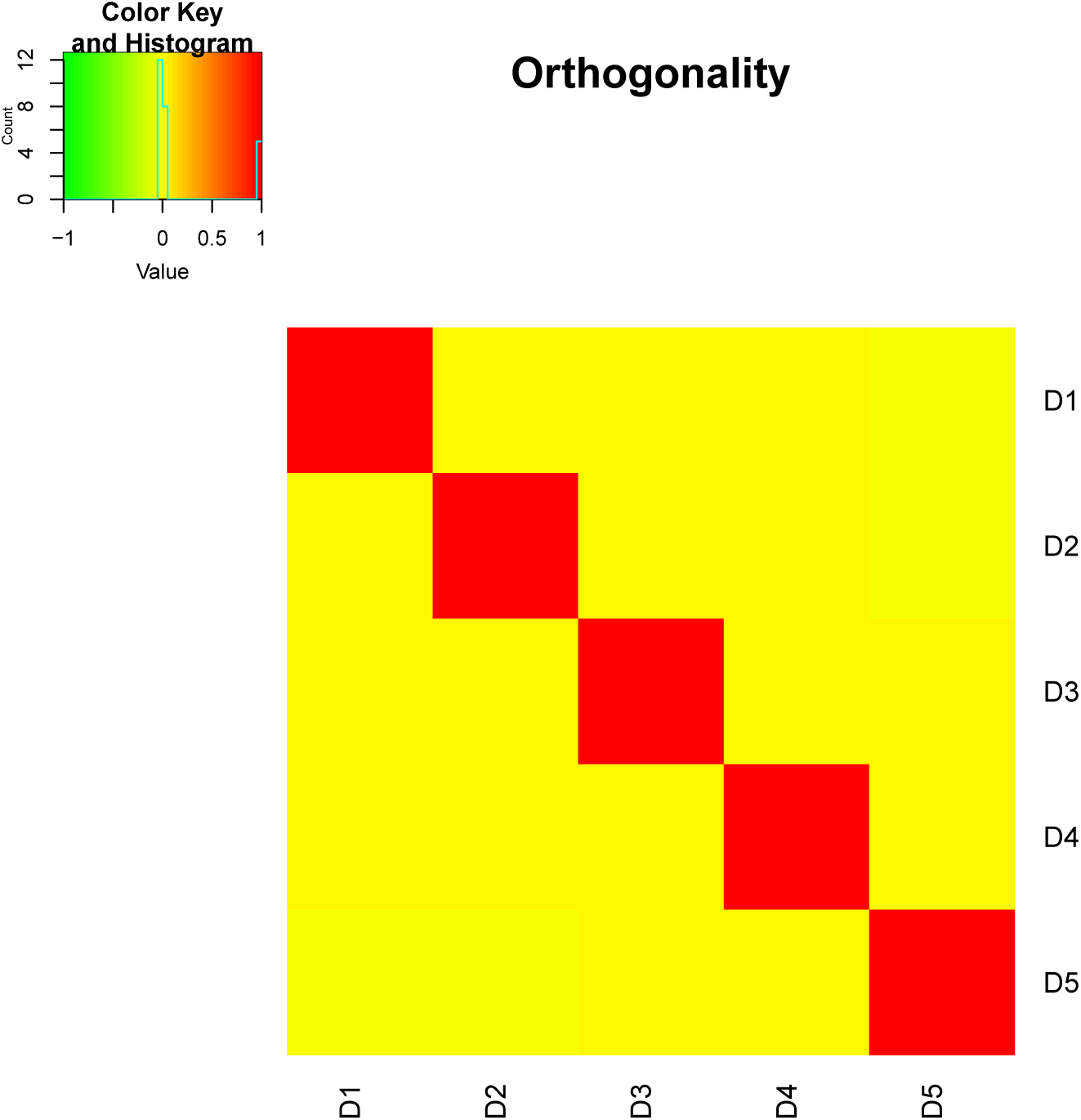
Orthogonality of the five identified projection directions from CAP.

**Table F.1:** Similarity between the five projecting directions from CAP and the seven PCs from CAP-C method.

|  |  | CAP-C |  |  |  |  |  |  |
| --- | --- | --- | --- | --- | --- | --- | --- | --- |
|  |  | CPC4 | CPC6 | CPC8 | CPC10 | CPC17 | CPC18 | CPC19 |
|  | D1 | -0.115 | -0.065 | 0.020 | 0.140 | 0.024 | 0.202 | 0.165 |
|  | D2 | **0.638** | 0.144 | 0.053 | -0.051 | -0.086 | 0.139 | -0.078 |
| CAP | D3 | -0.193 | -0.311 | -0.007 | 0.056 | -0.067 | -0.118 | -0.096 |
|  | D4 | -0.072 | 0.051 | 0.208 | 0.234 | -0.329 | **-0.669** | -0.339 |
|  | D5 | -0.214 | 0.432 | 0.192 | -0.016 | 0.009 | -0.153 | 0.349 |

To study the reliability of our proposed method, we apply the same linear projection to the rest three sessions of resting-state fMRI data acquired from the same subjects in the HCP study. Figure F.11 shows the estimated model coefficients and 95% bootstrap confidence interval. From the figure, the estimate and significance are very similar to the result presented in Figure 3 of Section 5, which postulates the existence of difference between age groups and/or sex within these five subnetworks of the DMN.

**Figure F.11:**
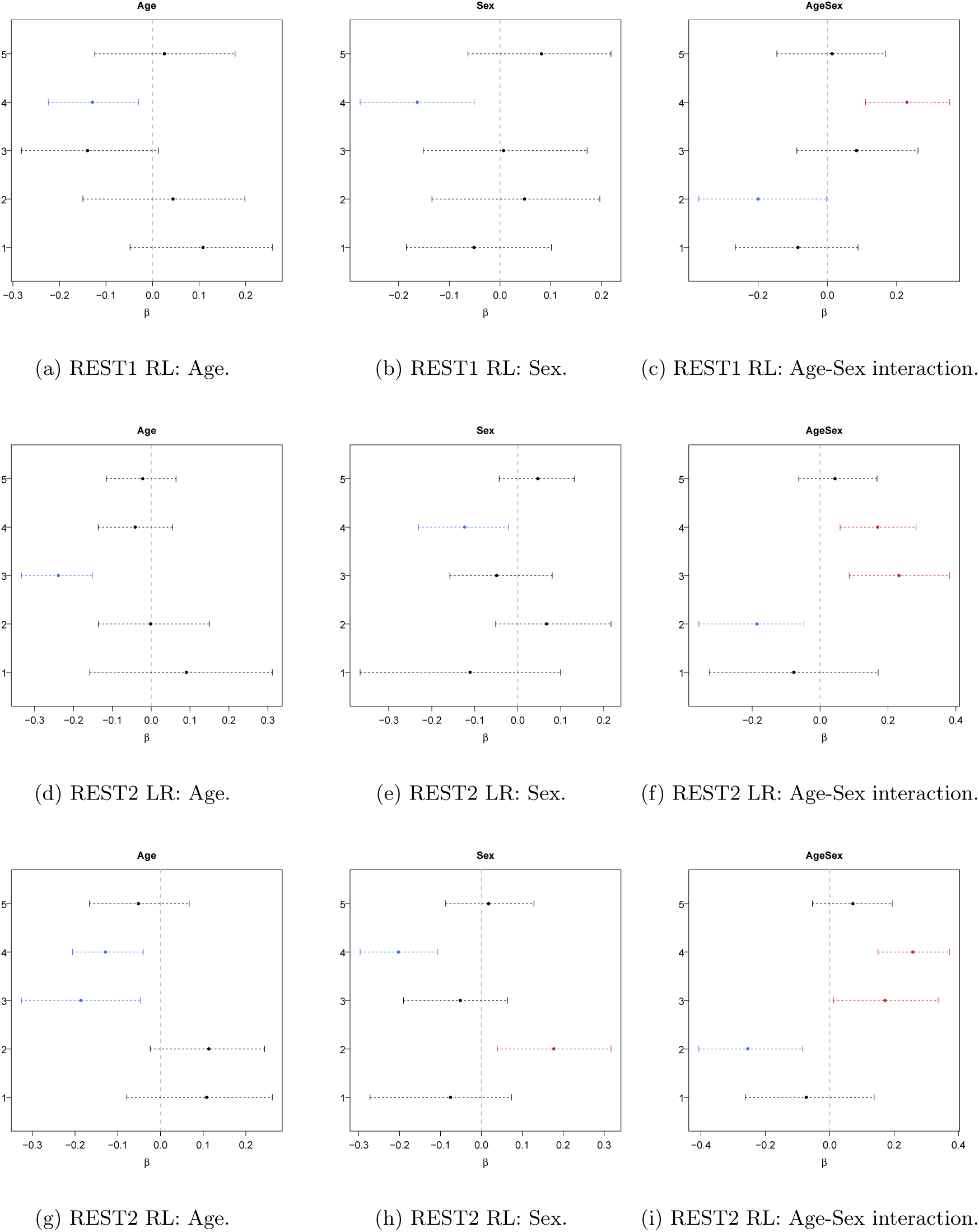
Estimated model coefficients and 95% bootstrap confidence interval of the five identified projection directions from the CAP approach in Section 5 tested on the rest three sessions of resting-state data collected from the same subjects in the HCP study.

**Figure F.12:**
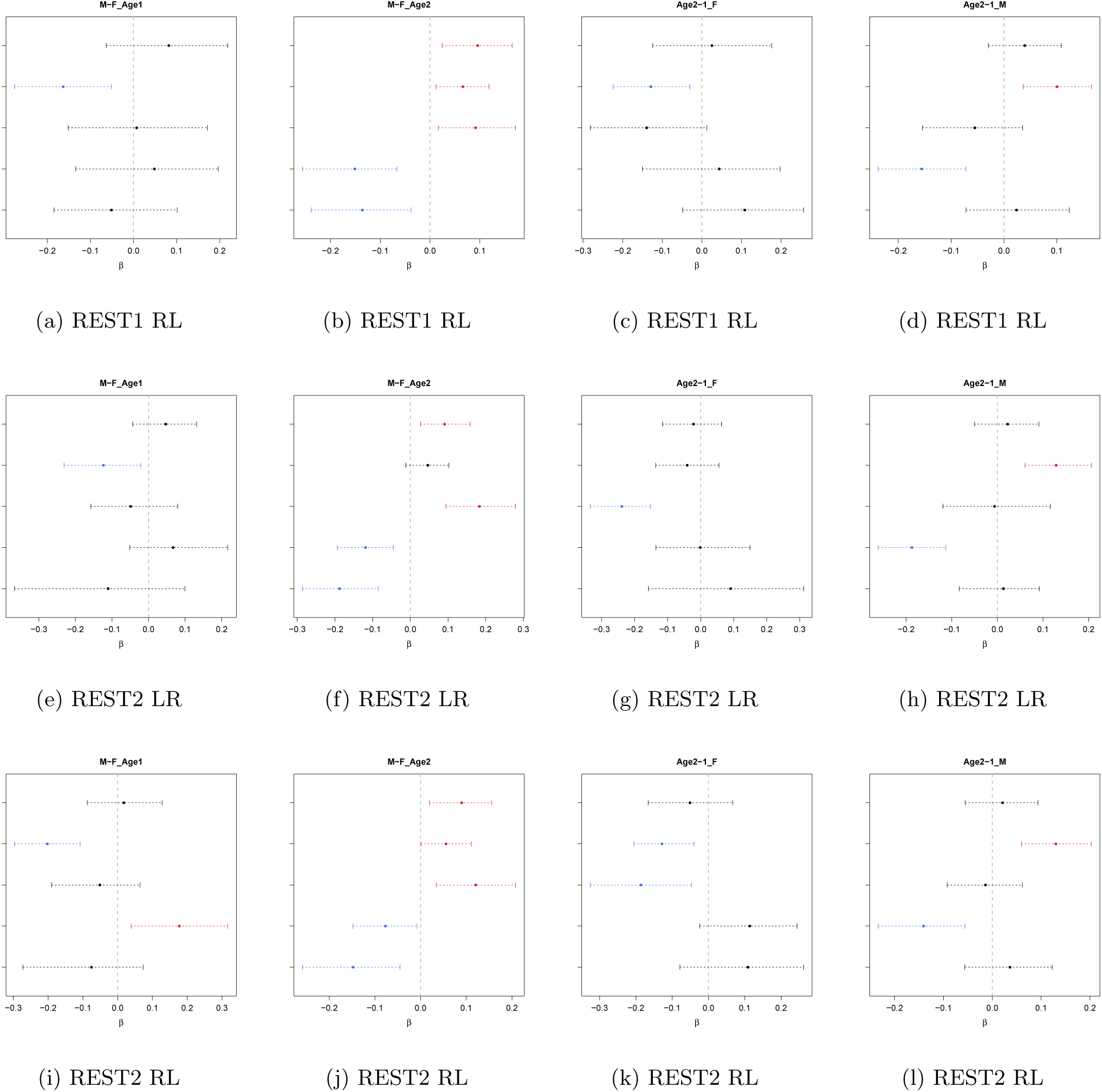
Pair-wise comparison of the five identified projection directions from the CAP approach in Section 5 tested on the rest three sessions of resting-state data collected from the same subjects in the HCP study.

